# Comparative genomics of *Dolichospermum circinale* strains with differential paralytic shellfish toxin profiles

**DOI:** 10.64898/2026.06.11.731795

**Authors:** Joao P. A. Pereyra, Paul M. D’Agostino, Verlaine J. Timms, Torsten Thomas, Brett A. Neilan

## Abstract

The cyanobacterium *Dolichospermum circinale* is a known producer of the neurotoxin saxitoxin and its analogues, collectively known as the paralytic shellfish toxins (PSTs). PSTs vary in potency, and the reported toxin profiles of *D. circinale* blooms vary in the quantities of individual PSTs, with the regulation of these profiles being poorly understood. In this study, we present the genomes of four *D. circinale* strains (ACBU01, ACMB03, ACMB13 and FSS-124) with unique PST profiles and perform genome-wide comparisons and specific analysis of the PST-producing biosynthetic gene cluster (*sxt*) to understand the variability in PST quotas. A reassessment of the previously published *D. circinale* AWQC131C genome was also performed to collate genomic variation between all strains. Analysis at the nucleotide and amino acid sequence level revealed that toxic strains maintain high genome-wide similarities, corroborated by the analysis of the pan- and variable genomes of each strain. Specifically, the *sxt* gene sequences were 99-100% identical across all strains. Novel tailoring (*sxtSUL*, *sxtDIOX*) and transport (*sxtM4*) genes were identified within the *sxt* cluster that were not reported previously in *D. circinale*. Taken together, these results indicate that the genetic machinery involved in PST production is conserved in this species, suggesting that the regulation of PST biosynthesis in *D. circinale* does not occur at the genomic level.

## Introduction

Paralytic shellfish toxins (PSTs) are potent neurotoxic alkaloids and the causative agents of paralytic shellfish poisoning. The toxicological effects of PSTs have been well documented, with human intoxication and fatalities reported over the years (Etheridge, 2010). The incidence of PST-producing blooms has increased worldwide due to nutrient enrichment of waters and rising temperatures brought on by climate change (Pearson et al., 2016, Paerl and Otten, 2013). The identification and characterisation of PST-producing strains for the protection of human health is vital in the face of these increasing ecological risks, especially given the vast difference in the potency of individual PSTs.

PSTs are tricyclic compounds (trialkyl tetrahydropurines) consisting of more than 57 natural derivatives, of which saxitoxin is regarded as the parent molecule (Wiese et al., 2010). These analogues differ structurally at four positions, where they may be non-sulfated (such as saxitoxin and the N1-hydroxylated neosaxitoxin), mono-sulfated (such as the gonyautoxins; GTXs), di-sulfated (C-toxins) or lack a carbamoyl moiety (dc-toxins) (Wiese et al., 2010). Non-sulfated PSTs are the most potent, followed by mono-sulfated and then di-sulfated PSTs (Wiese et al., 2010).

Cyanobacterial harmful algal blooms (cHABs) vary in their toxicity and toxin profile, which is attributed to the worldwide geographical differentiation of cyanotoxin-producing strains. PST production has been reported in several cyanobacterial species such as *Aphanizomenon gracile* (found in Norway, Spain, Germany and the USA), *Microseira wollei* (USA), *Raphidiopsis raciborskii* (Brazil), *Scytonema crispum* (New Zealand) and *Dolichospermum circinale* (Australia), (Ballot et al., 2016, Humpage et al., 1994, Carmichael et al., 1997, Soto-Liebe et al., 2010, Smith et al., 2011). In Australia, the PST profiles of toxic *D. circinale* strains are commonly dominated by di-sulfated PSTs C1 and C2, which constitute approximately 90% of the toxin quota, with the remaining 10% consisting of the various non-N1 hydroxylated PSTs (Velzeboer et al., 2000, Llewellyn et al., 2001, Pereyra et al., 2017). However, there is significant variation in the quota of the more potent non-disulfated PSTs produced by these strains. The composition of the PST profile and the intracellular to extracellular toxin ratio in *D. circinale* have been shown to be affected by environmental factors such as nitrate concentration, pH and salinity (Velzeboer et al., 2001, Ongley et al., 2016), suggesting the presence of regulatory genetic elements that respond to these factors. Additionally, a comparison of toxic and non-toxic *D. circinale* strains demonstrated differences in proteomic responses to external stimuli (phosphate depletion and increased extracellular NaCl) (D’Agostino et al., 2016).

Crucially, the genetic element that confers PST toxicity is the presence of the *sxt* gene cluster, initially detected in *Raphidiopsis raciborskii* T3 (Kellmann et al., 2008a). Since then, homologous clusters have been identified in *D. circinale* AWQC131C (hereafter AWQC131C), *Aphanizomenon gracile* NH-5 (Mihali et al., 2009), *R. brookii* D9 (Stucken et al., 2010), *M. wollei* (Mihali et al., 2011), and most recently in *A. gracile* NIVA-CYA 851, NIVA-CYA 655, NIVA-CYA 676 and UAM 529 (Ballot et al., 2016), *S. crispum* CAWBG524 (= *S*. cf. *crispum* UCFS10) and CAWBG72 (= *S*. cf. *crispum* UCFS15) (Cullen et al., 2018).

While each *sxt* cluster appears to be species-specific, it is similar in the overall composition and organisation of genes within strains. The formation of saxitoxin analogues that make up unique PST profiles is attributed to specific tailoring enzymes within the *sxt* gene cluster, using saxitoxin as a precursor (Kellmann et al., 2008a). Several of these tailoring genes have recently been characterised *in vitro*, the first being the N-sulfotransferase SxtN (Cullen et al. 2018), followed by the C-11 hydroxylase SxtDIOX (Lukowski et al. 2018). Interestingly, the functional elucidation of the *sxtDIOX* gene in the formation of GTXs and C-toxins urges a reassessment of toxic *D. circinale* strains, given the absence of this gene in all reported *D. circinale sxt* clusters. Furthermore, there are currently no comparative genomic studies of PST-producing *D. circinale* strains, despite investigations in other strains correlating variation in the *sxt* gene cluster to differential PST profile formation (Cullen et al., 2018).

Here we reanalyse the genome of the reference strain (AWQC131C) and present the genomes of four additional toxic strains of *D. circinale* (ACBU01, ACMB03, ACMB13 and FSS-124), isolated from three distinct regions of Australia. A survey of these four strains revealed unique PST profiles in all isolates, with the difference in individual and total PST quota not being explained by cellular morphology, phylogeny or geography (Pereyra et al., 2017). A whole-genome sequencing approach is employed to assess the variability and/or similarity of genetic elements involved in PST production and regulation in toxic *D. circinale* strains, providing insight into the geographic homogeneity of Australian PST producers. The identification of factors affecting PST production is an important step in the management of cHABs and molecular monitoring of potentially toxic bloom events.

## Material and methods

### Bacterial strains, genomic DNA extraction and genome sequencing

Four toxic *D. circinale* strains from three unique locations around Australia were obtained from the CSIRO Australian National Algae Culture Collection: ACBU01 (from the Burrinjuk Dam, NSW), ACMB03 and ACMB13 (Mount Bold Reservoir, SA), and FSS-124 (Gordonbrook Dam, QLD). The strains were maintained in MLA medium (Bolch and Blackburn, 1996) at 20 °C with 30 to 80 µmol photons m^-2^ s^-1^ at a 12:12 hour light:dark cycle.

Genomic DNA was extracted from all four strains as described by Morin et al. (2010), except for an additional washing step. Specifically, the cyanobacterial filaments were filtered onto a Whatman GF/C 1.7 µm glass microfiber filter and washed with two volumes of fresh MLA media. Cells were freeze-thawed to damage the cell wall, enzymatically lysed with 50 mg/mL of lysozyme then treated with 50 mg/mL of proteinase K in 2% sodium dodecyl sulphate (SDS) to degrade proteins. The lysate was treated with 1% cetyl trimethylammonium bromide (CTAB) solution in the presence of NaCl (with a final concentration of 1M) to remove polysaccharides, proteins and cell wall debris. Chloroform-isoamyl alcohol was then used for nucleic acid extraction. DNA was precipitated with isopropanol then washed with ice-cold 70% ethanol. TE buffer was used for the final resuspension of the DNA pellet before treatment with 10 mg/mL of RNAse. Genomic DNA concentration and quality was measured via Qubit fluorometric quantitation (Thermo Fisher Scientific) and agarose gel electrophoresis.

Genomic libraries were prepared using the Nextera XT DNA Library Preparation Kit, and 2 × 300 bp sequencing was performed on the Illumina MiSeq platform at the Ramaciotti Centre for Genomics at the University of New South Wales.

### Genomic assembly and binning

Paired sequencing reads were quality processed by removing Illumina adapters in palindrome mode. Quality processing of the bases was then performed by removing bases at the start and end of the reads, if the quality was below 3. Bases were also removed if four continuous bases had an average quality less than 15. After quality processing, only reads of length 36 bp or greater were kept. This entire process was performed with Trimmomatic (Bolger et al., 2014) version 0.38. The quality of the reads was assessed using FastQC software (https://www.bioinformatics.babraham.ac.uk/projects/fastqc/) version 0.11.5.

Processed reads were assembled using SPAdes (Bankevich et al., 2012) version 3.12.0, using k-mer sizes 21, 33, 55, 77, 99 and 127. Coverage information was obtained by mapping the reads of each genome to the contigs using Bowtie2 (Langmead and Salzberg, 2012) and sorted with Samtools (Li et al., 2009). As the cultures were not axenic, contig binning was performed using MetaBAT (Kang et al., 2015) with default parameters. The quality of bins was assessed using CheckM (Parks et al., 2015) to determine genome completeness, contamination and marker lineage for each bin.

### Taxonomic and phylogenetic analysis

The four cyanobacterial genomes were uploaded to the Microbial Genome Atlas (MiGA) (Rodriguez et al., 2018) to identify the ten most closely related strains from the NCBI RefSeq database considering the average amino acid identity (AAI). The non-cyanobacterial genomes produced by the contig binning for all strains were also uploaded to MiGA for AAI-based taxonomic identification of possible cyanotoxin-degrading bacteria and, where possible, 16S rRNA genes. In addition to the ten closest genomes, the reference genomes of the PST-producing AWQC131C, *S. crispum* CAWBG524 (=UCFS10) and *S. crispum* CAWBG72 (=UCFS15) and the non-toxic *D. circinale* AWQC310F (hereafter AWQC310F) were included to calculate genomic similarity via average nucleotide identity (ANI). The genomes of *Agrobacterium fabrum* str. C58, *Escherichia coli* str. K12 substr. MG1655 and *Gloeobacter violaceus* PCC 7421 were used as outgroups. The OrthoANIu algorithm was used to calculate pairwise ANI between all the genomes (Yoon et al., 2017). The comparative genome analysis based on AAI was repeated, this time including all genomes used for ANI calculation, using CompareM (https://github.com/dparks1134/CompareM).

For the phylogenetic analysis of the genomes, a specific set of 198 marker protein sequences (Supplementary Material Table S1) were identified from all bacterial genomes in the NCBI RefSeq database. These marker proteins were present in ≥95% of genomes in the database and existed as single-copy genes in ≥95% of cases, facilitating the comparison of phylogenetically diverse genomes (including non-cyanobacterial outgroups). A profile hidden Markov model (hmm) was generated for the 198 marker proteins using HMMER (Eddy, 1998). Using the same genomes as for the ANI and AAI calculation, the predicted genes of each genome were searched using HMMER for hits against the 198 hmm profile and subsequently concatenated. The concatenated protein sequences were aligned using MAFFT (Katoh and Standley, 2013). A phylogenetic tree was constructed with MEGAX (Kumar et al., 2018) using the maximum likelihood method with 1000 bootstraps and the Jones-Taylor-Thornton substitution model.

### Genomic annotation and comparative analysis of genomes

All four newly-sequenced genomes were submitted to the Integrated Microbial Genomes and Microbiomes system (IMG/M) version 5.0 and processed through the IMG annotation pipeline (Chen et al., 2019). The nucleotide sequence of each genome is publicly available from the US DOE JGI server under the Taxon ID 2821308615, 2821300090, 2821304336 and 2821292572 for ACBU01, ACMB03 and ACMB13 and FSS-124, respectively. The reference AWQC131C (Taxon ID 2516653039) and AWQC310F (Taxon ID 2516653040) genomes were found in the expert review system IMG/MER (as well as the NCBI database) and were included in parts of the comparative analysis.

The comparative analysis of genomes was performed based on the results of the annotation and the subsequent classification of protein coding genes via the KEGG (Kanehisa and Goto, 2000), MetaCyc (Caspi et al., 2018), COG (Tatusov et al., 2000), Pfam (El-Gebali et al., 2019) and TIGRFAM (Haft et al., 2001) databases. Pathways within the genomes were manually searched using Metacyc and UniProt (Bateman et al., 2019). The IMG/MER system was used for inferring synteny between the genomes using MUMmer (Kurtz et al., 2004) and for genome clustering based on functional profiles.

Identification of non-ribosomal peptide synthetase (NRPS) and polyketide synthase (PKS) domains was performed using the antiSMASH software version 5.0 (Blin et al., 2019). The software was run using multiple detection strictness parameters: strict, relaxed and loose. The latter was used for a more flexible search of cryptic clusters that are less well-defined.

### Sequence analysis of the *sxt* gene cluster

The *sxt* gene cluster was curated and analysed manually using both the UniPro UGENE software toolkit (Okonechnikov et al., 2012) version 1.32.0 and the Geneious Software Suite (Kearse et al., 2012) version 5-6.0.5. Nucleotide sequences obtained from Illumina genome sequencing, PCR products and homology searches from the NCBI database were aligned and visualised using these software packages. Sequence similarity of *sxt* genes between all four genomes and the reference AWQC131C genome was assessed by performing a Basic Local Alignment Search Tool (BLAST) at the nucleotide (BLASTn) and the protein (BLASTp) levels using default parameters. The nucleotide spacing and sequences between each gene were also compared via BLASTn. The regions upstream of open reading frames (ORFs) within the cluster were also analysed for potential variation in putative promoter sequences.

### PCR of DNA fragments

To confirm the arrangement of *sxt* genes in the four strains in this study, primers were designed for the amplification of regions of the *sxt* gene cluster in ACBU01, ACMB03, ACMB13 and FSS-124 (Supplementary Material Table S2). PCR was performed using KAPA HiFi HotStart ReadyMix (Roche), in a 20 µL reaction volume as per the manufacturers’ instructions with a final concentration of 0.05 µM of each primer. Cycling conditions had an initial denaturation step at 95 °C for 3 min, followed by 30 cycles of DNA denaturation at 98 °C for 20 s, primer annealing at 58 °C for 15 s, DNA amplification at 72 °C for 1 min/kbp amplified, and a final extension at 72 °C for 10 min (Eppendorf Mastercycler 5333 PCR Thermocycler). The size and concentration of PCR amplicons were checked via agarose gel electrophoresis.

## Results

### Genomic structure of assembled *D. circinale* genomes

For all four strains the genomic binning produced a single bin with a cyanobacterial lineage assignment. The quality assessment of these four bins indicated high-quality with over 99% completeness with no contamination. Each bin also contained one 16S rRNA gene sequence that could be assigned to the species *D. circinale.* The genome assemblies of *D. circinale* strains ACBU01, ACMB03 and ACMB13 produced very similar metrics, with slight variation from FSS-124 in several statistics such as size, N50 and gene count (Table 1). The total assembly size for each genome was 4.4 Mbp for strains ACBU01, ACMB03 and ACMB13 and 4.6 Mbp for FSS-124. The FSS-124 strain had a slightly higher gene count (4,517) and the CDS count (4,440). However, all four strains were very similar in GC content (approximately 37.3%) and had the same proportion of *sxt* genes making up the total CDS for each genome (approximately 1%).

**Table 1:** Genome assembly statistics of *D. circinale* strains.

| Genome Characteristics | ACBU01 | ACMB03 | ACMB13 | FSS-124 |
| --- | --- | --- | --- | --- |
| Size (Mbp) | 4.4 | 4.4 | 4.4 | 4.6 |
| GC content (%) | 37.3 | 37.3 | 37.3 | 37.2 |
| N50 (bp) | 120,937 | 114,207 | 120,937 | 165,589 |
| Scaffold count | 57 | 61 | 60 | 46 |
| Shortest scaffold (bp) | 3919 | 3562 | 3919 | 3501 |
| Longest scaffold (bp) | 299,977 | 331,811 | 300,234 | 315,827 |
| Avg. scaffold length (bp) | 77,576.4 | 72,502.4 | 73,806.3 | 100,465.3 |
| Gene count | 4,284 | 4,245 | 4,278 | 4,517 |
| Coding Base Count NP % | 82.73 | 82.55 | 82.69 | 82.95 |
| CDS count | 4,206 | 4,167 | 4,200 | 4,440 |
| rRNA genes | 8 | 8 | 8 | 6 |
| tRNA genes | 41 | 41 | 41 | 41 |
| Genes with functional annotations (%) | 67.81 | 68.27 | 67.88 | 66.15 |

### Taxonomic and phylogenetic analysis

Comparative genomic analysis identified the ten most closely related species for all four genomes as: *Dolichospermum compactum* NIES 806 (NCBI Accession NZ_AP018316.1), *Anabaena* sp. WA102 (NC_007413.1), *Anabaena* sp. 90 (CP003284.1 and CP003285.1), *Nostoc azollae* 0708 (NC_014248.1), *Cylindrospermum* sp. NIES 4074 (AP018269.1), *Nostoc carneum* NIES 2107 (AP018180.1), *Aulosira laxa* NIES 50 (NZ_AP018307.1), *Anabaena variabilis* ATCC 29413 (NC_007413.1), *Nostoc sphaeroides* (strain Kützing, NZ_CP031941.1) and *Anabaenopsis circularis* NIES 21 (NZ_AP018174.1). A subsequent ANI analysis including reference genomes of known PST-producers revealed that all four *D. circinale* genomes had the closest genomic similarity to each other and the AWQC131C (>98%) and AWQC310F (>96%) genomes (Supplementary Materials Figure S1). This was corroborated by identical results from the comparative analysis based on AAI (Supplementary Materials Figure S2).

AAI and 16S rRNA taxonomic analysis of non-cyanobacterial sequences found in the cultures demonstrated the absence of any genera known to degrade cyanotoxins, including those from the phyla Actinobacteria, Firmicutes, α-, β- and ɣ-Proteobacteria (Kormas and Lymperopoulou, 2013).

The phylogenomic analysis of the genomes demonstrated the close relationship between the four *D. circinale* strains in this study and the reference AWQC131C and AWQC310F strains, which formed their own clade separate from the other *Anabaena* and *Dolichospermum* species (Figure 1). Within this clade, the four genomes grouped closer to the toxic AWQC131C than the non-toxic AWQC310F. These results supported the taxonomic analysis of the strains and suggested genome-wide similarities between toxic *D. circinale* strains.

**Figure 1:**
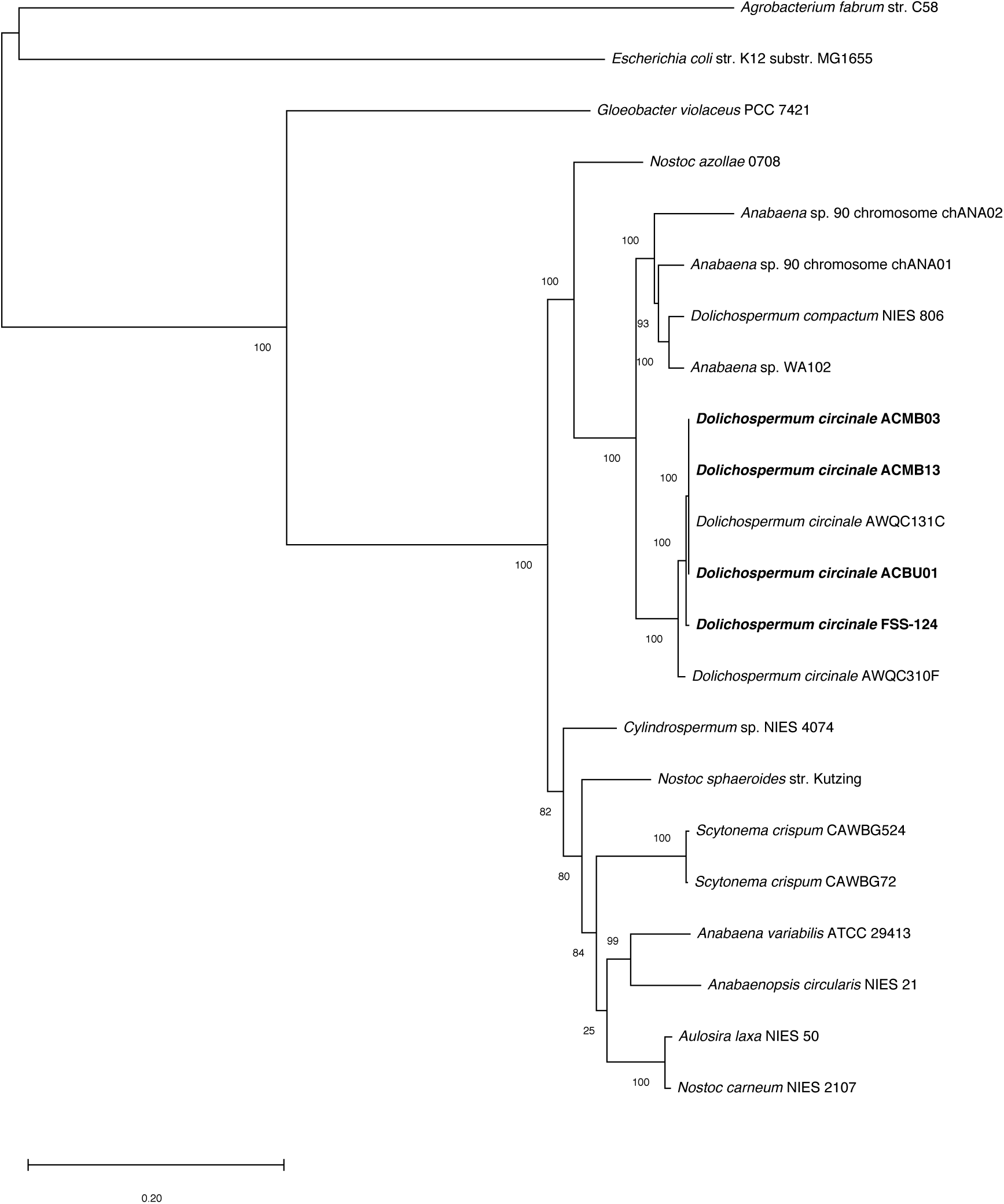
Phylogenomic tree of *Dolichospermum* and related species based on 198 concatenated protein sequences. The four genomes from this study are in bold type.

### Comparative analysis of genomes

The comparison of identified COGs in each strain revealed 86 COGs unique to FSS-124 and 48 COGs present in ACBU01, AMCB03 and ACMB13 only (Figure 2; Supplementary Material Table S3). FSS-124 had several unique COGs of unknown or general function predictions, as well as elements involved in the mobileome (i.e. prophages and transposons) and post-translational modification, protein turnover and chaperones. In contrast, the other three strains contained unique COGs involved in carbohydrate transport and metabolism, as well as cell motility. Comparison of the four genomes with the reference AWQC131C genome gave similar results, with FSS-124 showing the greatest amount of variation between all toxic strains.

**Figure 2:**
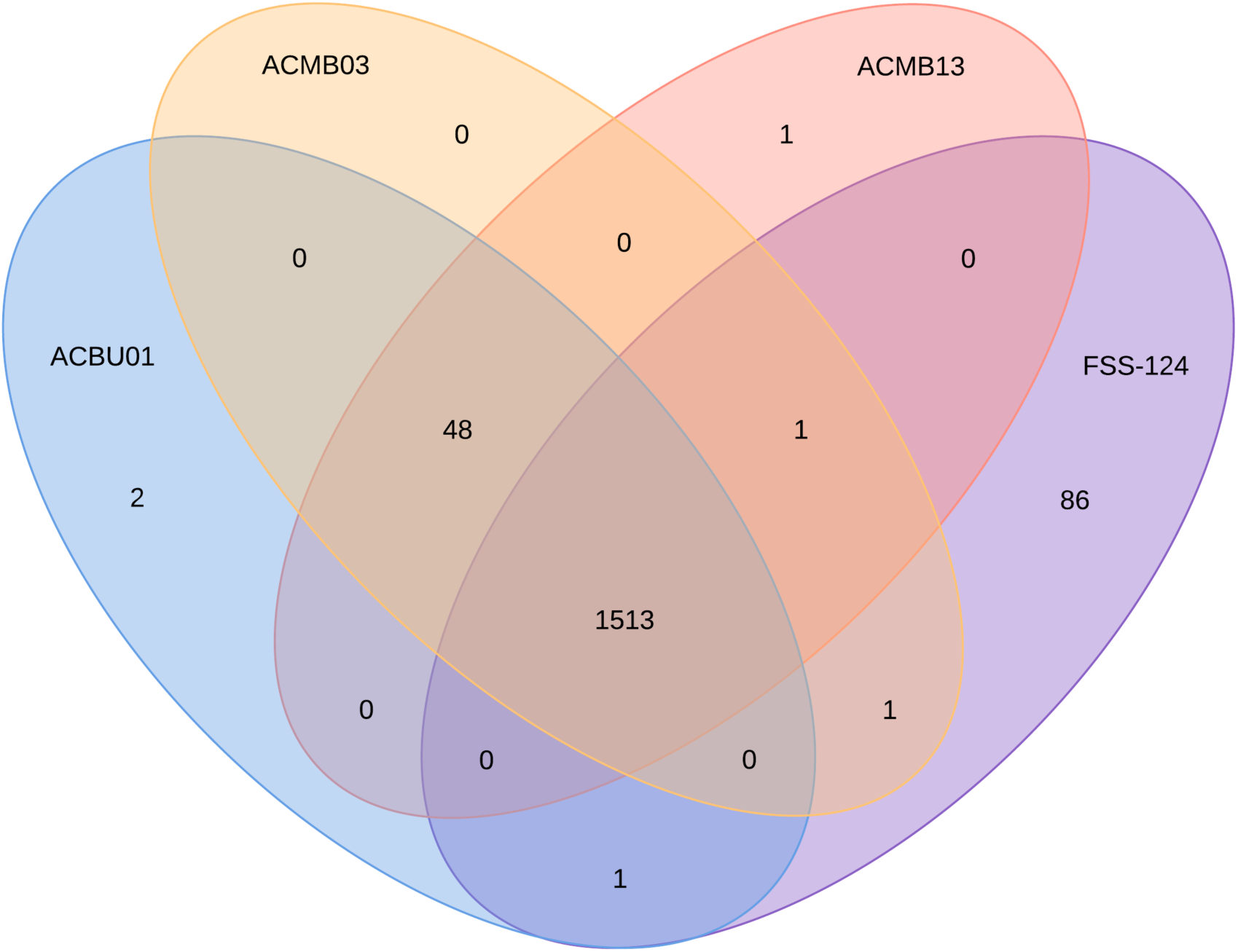
COG distribution across the four *D. circinale* genomes based on IMG annotation. All values indicate the number of COGs present in one or more genomes.

Other annotations also suggested a slight variation between FSS-124 and the other three strains (Supplementary Material Tables S4-S6), with the numbers of unique genes based on KEGG Orthology (21 compared to 18), Pfam (60 compared to 40) and TIGRfam (24 compared to 11) corroborating with the analysis based on COGs. FSS-124 had more protein-coding genes involved in Lipid A biosynthesis, whereas the other three strains had more protein coding genes involved in fatty acid biosynthesis and cell metabolism pathways (such as gluconeogenesis, glycolysis and the pentose phosphate pathway). An analysis exploring the conservation of synteny across all four strains in this study again showed a marked difference between the FSS-124 genome and the genomes of the other three strains (Supplementary Material Figure S3).

A survey of the four genomes for regulatory elements such as sigma factors, sigma factor regulators, histidine kinases and serine/threonine kinases revealed minimal differences in the number and types of these elements. This remained consistent when comparing the four genomes to the reference AWQC131C (PST+) and AWQC310F (PST-) genomes, with the only exception being the total number of serine/threonine kinase genes between the four genomes (with a mean of 27 genes) and the reference genomes (with 23 genes each), however this was mostly due to gene duplications. Interestingly, the comparison of these genes at the amino acid level demonstrated that these genes were 100% identical in ACBU01, ACMB03 and ACMB13, but not always when compared to FSS-124 (97-100%), AWQC131C (99-100%) and AWQC310F (85-100%).

The analysis of transporters and antiporters showed the same trend, with the four genomes having more genes (with a mean of 145 transporters and 15 antiporters) than the reference genomes (with a mean of 132 transporters and 12 antiporters) due to gene duplications, maintaining the same variation in amino acid similarity. However, there were several transporter genes unique to the four genomes, including ATP-binding cassette (ABC) transporter genes for lipopolysaccharides, multidrug resistance, sugar and urea. Interestingly, a nickel transport protein was present only in toxic *D. circinale* strains, suggesting a possible regulatory mechanism specific to PST-producing strains.

The presence of nitrogen and phosphorus metabolism genes was conserved across all four strains. Among these were several genes with the potential to regulate the *sxt* gene cluster (present in the four genomes and AWQC131C), and therefore PST production, in response to environmental cues (Table 2). This includes the *pho* regulon with the histidine kinase *PhoR* and the transcriptional regulator *PhoB*, which were conserved across all *D. circinale* strains. The global nitrogen regulator NtcA and the ferric uptake regulators (Fur) were also identified in all *D. circinale* genomes. The only differences were the presence of gene for putative phosphate-binding proteins (PBPs) and alkaline phosphatase precursors (PhoA) in AWQC310F, but given that the strain is non-toxic, these genes are unlikely to contribute to affect PST variability. Variation between these proteins at the amino acid level was again evident (Table 2). Other annotated metabolic pathways were conserved across *D. circinale* genomes, with more than 99% of the pathways identified present in all four genomes of this study, and more than 97% when including the reference strains.

**Table 2:**
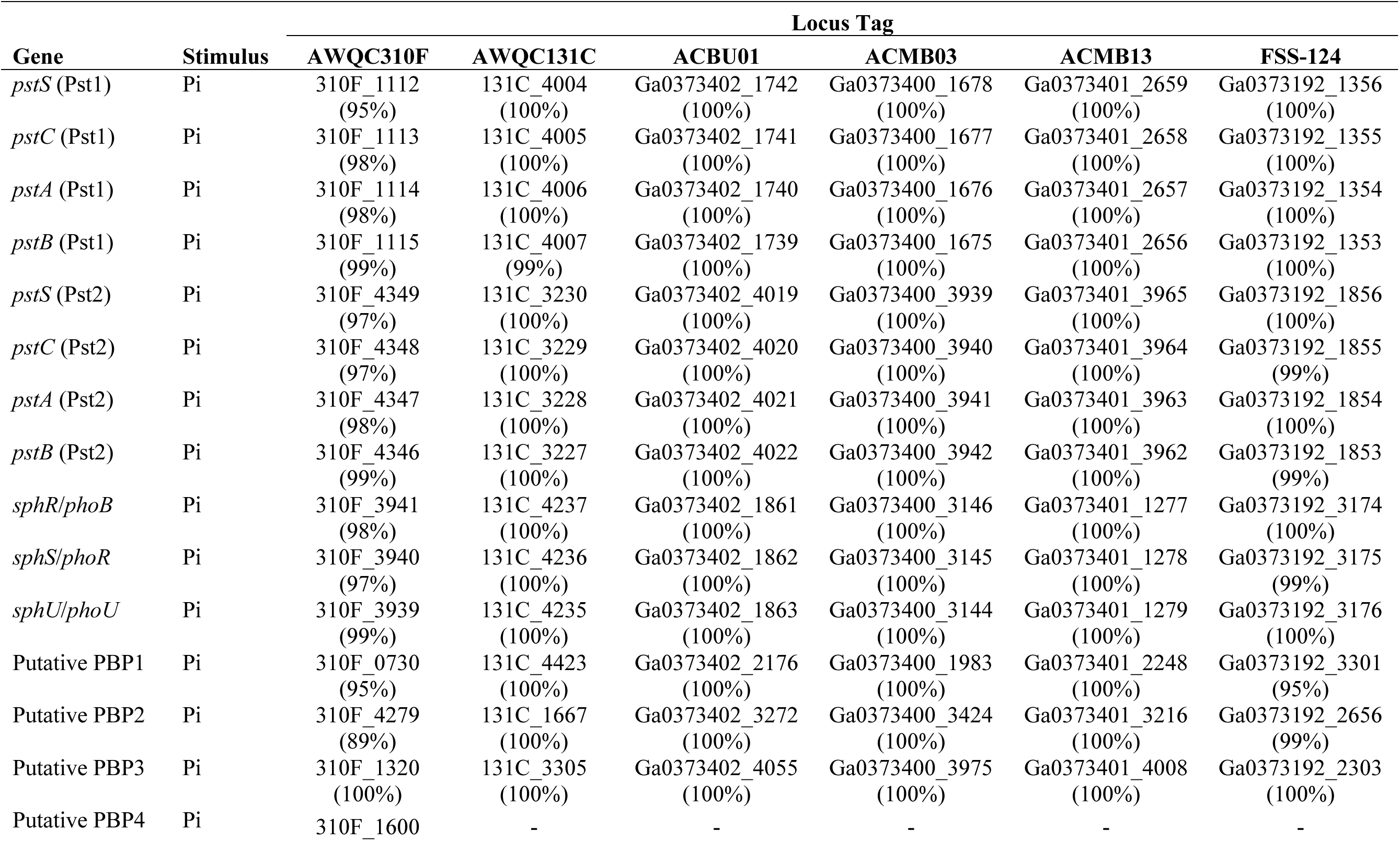
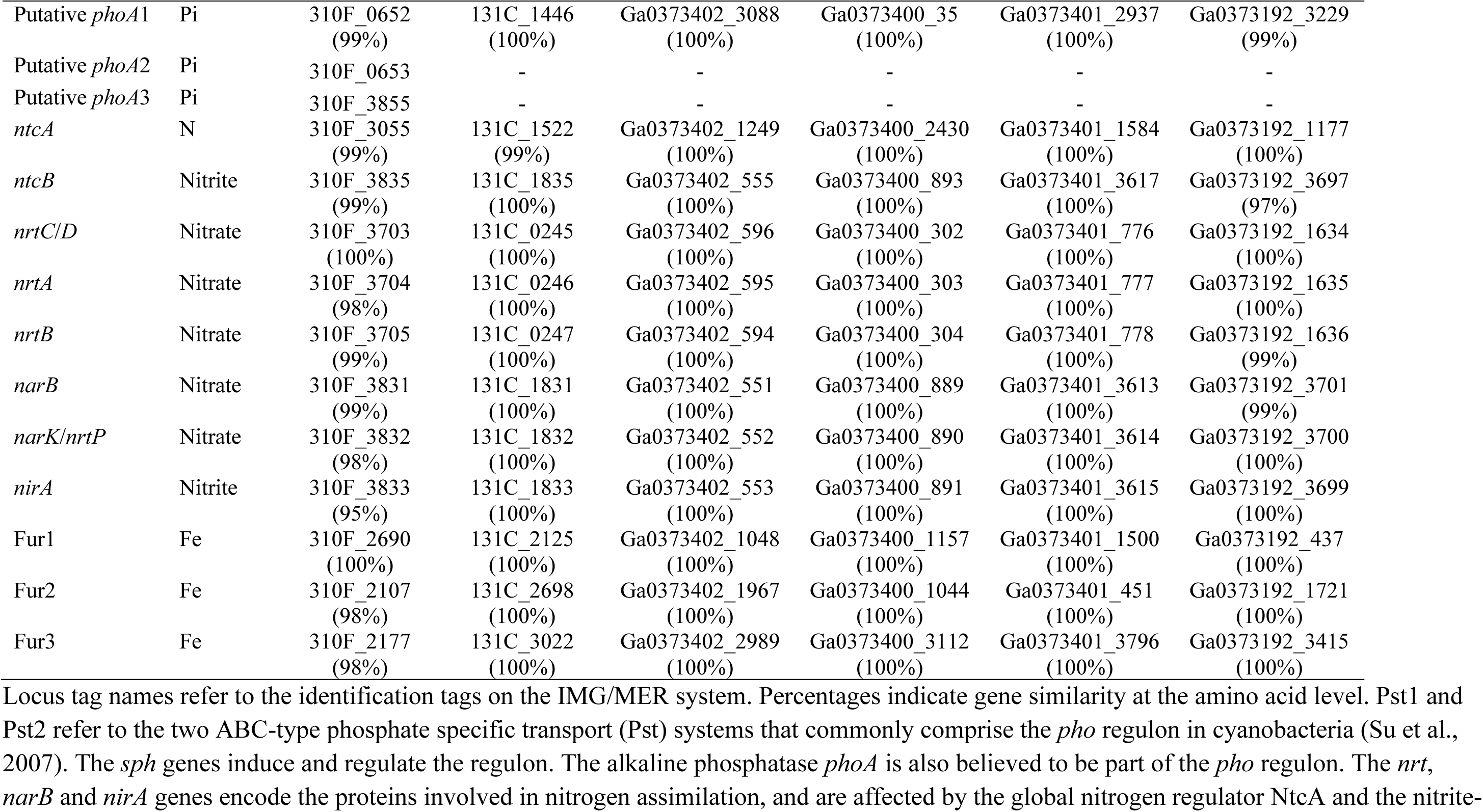
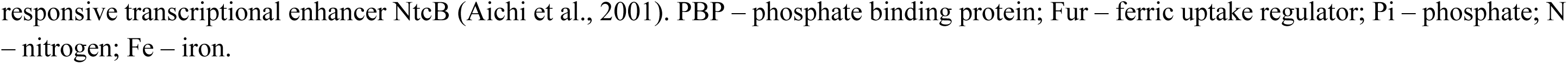
Putative regulatory elements/operons of the *sxt* gene cluster in *D. circinale*.

In addition to the *sxt* gene cluster (see below), three putative secondary metabolite biosynthesis gene clusters (from a total of 32-33 cryptic clusters) were identified in the four *D. circinale* genomes sequenced in this study and in the AWQC131C and AWQC310F reference genomes (Supplementary Material Figures S4-S6). Results from an antiSMASH analysis were filtered based on a >80% similarity threshold of the identified clusters to known natural product clusters. The cluster with the highest similarity (100%) was geosmin synthase, responsible for producing the bicyclic alcohol geosmin. The second identified cluster was for the ribosomally synthesised and post-translationally modified peptide (RiPP) anacyclamide (85% similarity), a cyanobactin common to the genus *Anabaena*. The third identified cluster was homologous to the *hgI* cluster (85% similarity), responsible for heterocyst glycolipid biosynthesis, containing additional polyketide synthase (PKS) modules downstream.

### Analysis of the *sxt* gene cluster

The *sxt* gene cluster was identified in all four *D. circinale* genomes and manually curated. The entire cluster was assembled from multiple scaffolds: two scaffolds for ACBU01 and ACMB13, and four scaffolds for FSS-124 and ACMB03. The expected *sxt* genes, based on the original AWQC131C cluster, were found in all strains, although further sequencing is required to confirm the sequence of five genes (*sxtT* and *sxtH* in ACMB03 and FSS-124, and *sxtM* in ACMB03) that had their sequence interrupted due to read cutoffs. Overall, the *sxt* gene clusters were identical across all four strains in terms of the presence, size and organisation of *sxt* genes. The *sxt* clusters were flanked by a β-lactamase gene at the 5′-end and a *smf* gene homolog at the 3′-end (Figure 3). A total of 30 ORFs were identified, including 28 *sxt* genes and one gene (ORF24) previously described in several *sxt* clusters including AWQC131C. This contrasted with the originally reported AWQC131C cluster that had only 23 genes (Mihali et al., 2009). Two of the ORFs, *sxtV1* and *sxtV2*, were present in all clusters and denoted a disrupted *sxtV* pseudogene, also present in AWQC131C.

**Figure 3:**
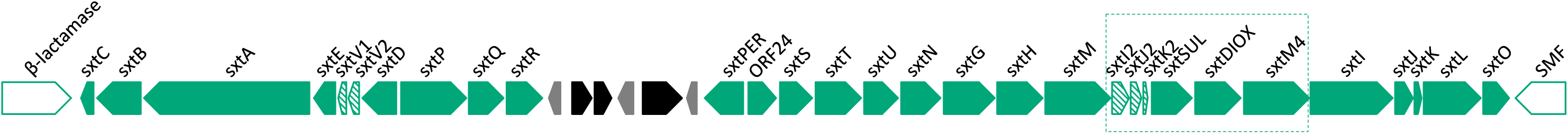
The *sxt* gene cluster found in the toxic *D. circinale* genomes sequenced in this study. The dotted box represents new genes incorporated into the amended cluster. The genes flanking the cluster are shown in white. Truncated genes are shown with diagonal lines. ORFs unrelated to PST biosynthesis are shown in grey and black (putative transposases).

The consensus *sxt* cluster from the genomes sequenced in this study had several additional (and hitherto novel) ORFs, compared to the *sxt* cluster originally described from AWQC131C, including one full-length gene (*sxtDIOX*), three truncated genes (*sxtI2*, *sxtJ2* and *sxtK2*) and one paralog (*sxtM4*). Additionally, the *sxtSUL* gene was identified within the consensus cluster between the *sxtM* and *sxtI* (Figure 3). Confirmation of the new *sxt* genomic arrangement was performed via PCR, using primers flanking *sxtSUL* and *sxtI*, *sxtSUL* and the *smf* gene, and *sxtG* and *sxtI*, yielding amplicons of the expected sizes (Supplementary Material Figures S7-S9).

The *sxtSUL* (N-sulfotransferase) and *sxtDIOX* (C-11 hydroxylase) genes aligned most closely with their homologues in *S. crispum* CAWBG72. Additionally, the *sxtM4* paralog (a putative multi-drug and toxic compound extrusion (MATE) protein) was more closely related to the *sxtM* of *A. gracile* NH-5 at both the nucleotide and protein level. The sequences of *sxtI2*, *sxtJ2* and *sxtK2* were much shorter and revealed excisions when compared to their full-length homologs (putative O-carbamoyltransferase genes hypothesised to work in conjunction) in AWQC131C.

These additional sequences found in the four genomes of this study were also present in AWQC131C. The region between *sxtM1* and *sxtM4* was previously sequenced (D’Agostino, 2013) and the presence, size and organisation of the secondary ‘*sxt2*’ genes were identical to the additional sequence found in the *sxt* clusters of the four genomes sequenced of this study. This indicates that the genomic location of the ‘*sxt2*’ genes were within the main *sxt* gene cluster, suggesting a conserved cluster across *D. circinale* (Figure 3).

There were several single nucleotide polymorphisms (SNPs) within the genes in the *sxt* cluster across all four strains when compared to their homologs in AWQC131C. The *sxtP* and *sxtDIOX* genes in FSS-124, as well as the *sxtM* and *sxtI* genes of all four strains in this study contained non-synonymous SNPs when compared to the corresponding genes found in AWQC131C. Further, the non-coding DNA sequence between genes (known as the intergenic spacer or IGS regions) of the *sxt* gene cluster showed no variation in terms of size and nucleotide sequence for ACBU01, ACMB03 and ACMB13 when compared to AWQC131C. However, there were SNPs found in the IGS between *sxtA* and *sxtE* and between *sxtPER* and ORF24 in strain FSS-124. Despite this, the regions containing the five promoter elements of the *sxt* cluster (PsxtD, PsxtP, PsxtPER1, PsxtPER2 and Porf24) described by D’Agostino et al., (2020) were conserved across all toxic strains.

## Discussion

Characterisation of the toxin profile of *D. circinale* cHABs has been reported in Australia since the early 1990’s (Bowling and Baker, 1996). *D. circinale* toxin profiles are commonly dominated by the C-toxins however the minor constituent PSTs can vary significantly from strain to strain (Velzeboer et al., 2000, Llewellyn et al., 2001, Pereyra et al., 2017). Specific toxin profiles are not correlated to morphology, phylogeny or geography, suggesting genetic differences between strains may be responsible for the observed toxin profiles (Pereyra et al., 2017). Here we investigated the genomes of four Australian PST-producing strains of *D. circinale* to assess their overall genomic variability as well as genetic variation within the *sxt* gene cluster.

The taxonomic and phylogenetic analyses established a close relationship between the four strains in this study at the genomic level (as well as with AWQC131C) and found no evidence for the presence of any known PST-degrading bacteria that could affect the variation in PST profiles. The tendency for toxic strains to be grouped together has been shown in other cyanobacteria within the *Anabaena*, *Dolichospermum* and *Aphanizomenon* (ADA) clade (Österholm et al., 2020) with a closer grouping associated with strains that produce the same secondary metabolites. The high degree of similarity of the toxic strains in this study is again evidenced by the high proportion of shared genes using various annotation databases. This demonstrates that a large proportion of the pan-genome of toxic *D. circinale* belongs to the core genome. Results suggest slightly higher conservation within the four strains (>91% COGs) compared to other PST producers, with the core genome in a study of toxic *C. raciborskii* strains comprising 86% of the pan-genome (Willis et al., 2018).

Despite the high degree of similarity, there were clear differences in the variable genome (i.e. strain-specific genes) of FSS-124 compared to the genomes of the other three strains (ACBU01, ACMB03 and ACMB13) and the reference strain (AWQC131C). However, these did not account for the variability in the PST profile of this strain (Table 3). Previous analyses of the genomes and proteomes of other cyanotoxin-producing strains of *Microcytis aeruginosa* (Alexova et al., 2011) and *Raphidiopsis raciborskii* (Sinha et al., 2014, Willis et al., 2018) demonstrated that the variable genome encodes gene for a variety of adaptation-related processes independent of cyanotoxin production and suggested that strains represent unique ecotypes adapted to survival in a particular environmental niche. The genetic differences between FSS-124 and the other toxic strains may be a result of adaptation given the unique prophages and transposons found in its genome, perhaps due to the difference in latitude and microclimate (Table 3). It is thus possible that FSS-124 as well as the other three strains are unique ecotypes, much like AWQC131C and AWQC310F (D’Agostino et al. 2014), though the variance in toxin quota of toxic *D. circinale* strains does not appear to be due to geography (Pereyra et al., 2017).

**Table 3:** Collection locations and PST profiles of toxic *D. circinale* strains.

| Strain | C1/C2 |  | dcSTX | GTX5 | STX |  | Reference |
| --- | --- | --- | --- | --- | --- | --- | --- |
| ACBU01 | 96.42 | 3.58 | - | - | - | - | Pereyra et al., 2017 |
| ACMB03 | 90.95 | 2.80 | 4.60 | 1.65 | - | - | Pereyra et al., 2017 |
| ACMB13 | 90.40 | 4.11 | 2.50 | 0.25 | 2.75 | - | Pereyra et al., 2017 |
| FSS-124 | 94.83 | 1.93 | 3.24 | - | - | - | Pereyra et al., 2017 |
| AWQC131C | 64 | 16 | 7 | 11 | - | 2 | Llewellyn et al., 2001 |

The presence of identically annotated regulatory elements in all strains (including sigma factors, sigma factor regulators, histidine kinases and serine/threonine kinases) implies uniformity in the metabolic regulation of toxic and non-toxic strains of *D. circinale*. The difference between FSS-124 and the other three strains in the analysis of transporters and antiporters was minor, though interestingly the identification of a nickel transport protein specific to toxic *D. circinale* strains suggests another possible variable in the effect of nutrient availability on PST production. The analysis of putative transcription factors (Table 2) was specifically based on the identification of putative binding sites in other *sxt* clusters (such as the *pho* regulon in the *R. raciborskii* T3 *sxt* cluster) or other cyanotoxin gene clusters (such as NtcA and ferric uptake regulators in the promoters of the *mcy* cluster forming microcystin) (Ginn et al., 2010, Martin-Luna et al., 2006, Alexova et al., 2011). Again, these were conserved across toxic strains, and differences between FSS-124 and the other three strains were minor, though any possible effect of the genomic variation in FSS-124 would require further investigation of the toxic *D. circinale* strains in response to stimuli implicated in modulating PST production (such as the availability of nitrogen and phosphorus, pH, and osmotic stress) via other omics approaches. The presence of three putative secondary metabolite clusters was consistent across the four strains and had been previously found in the reference AWQC131C and AWQC310F strains (Österholm et al., 2020), with no evidence in affecting PST profile formation.

The annotation of genes in the *sxt* gene cluster of *D. circinale* strains has been amended based on the results of this study (Figure 3). The identification of the *sxtSUL* gene was significant given that this gene was not identified in the original AWQC131C *sxt* cluster and previous investigations had only reported the presence of the gene in toxic *D. circinale* ACMB13 (Soto-Liebe et al., 2010, Pereyra et al., 2017). The discovery of the genes encoding SxtSUL and SxtDIOX in these clusters was anticipated given that the combined activity of these enzymes (the O-sulfotransferase and C11 hydroxylase, respectively) produce the mono-sulfated GTX3 (Lukowski et al., 2019), that makes up a small part of the PST profiles of toxic *D. circinale* strains and is essential to biosynthesise the C-toxins, which constitute most of the *D. circinale* PST profile (Velzeboer et al., 2000, Llewellyn et al., 2001, Pereyra et al., 2017).

The discovery of several other genes in the *sxt* cluster was unexpected. SxtM, a putative MATE protein (i.e. PST transporter), has been shown to retain a relaxed specificity typical of the NorM class of MATE proteins (Ongley et al., 2016). The duplication of this sodium-driven antiporter in the revised *sxt* cluster suggests a greater capacity for the export of PSTs across the cell membrane. Interestingly, a previous study suggested that variability within the substrate recognition motif of MATE proteins SxtF and SxtM in *R. raciborskii* T3 resulted in a different binding affinity of each transporter to different PSTs (Soto-Liebe et al., 2013). Given the newly found SxtM4 is a paralog of SxtM, the study of the specificity of this efflux protein may provide more insights into PST quotas in *D. circinale*. Further, the excisions in *sxtI2*, *sxtJ2* and *sxtK2*, as well as the absence of the putative *sxtI* catalytic site GPRALGRS (Kellmann et al., 2008b) in *sxtI2* suggests these genes are inactive despite being identified as ORFs with start and stop codons.

The insertion of the hitherto novel *sxt* genes between *sxtM* and *sxtI* suggests a change in the transcriptional organisation of the *D. circinale sxt* cluster. The active transcription of the putative operons containing all *sxt* tailoring genes (i.e. *sxtL*, *sxtN*, *sxtO*, *sxtSUL* and *sxtDIOX*) during a harmful bloom event would result in PST profiles dominated by C-toxins, given all the genes required for their biosynthesis are present. Results from this study show that the *sxt* cluster is conserved across toxic *D. circinale* strains and would thus explain the abundance of C-toxins in Australian isolates. Furthermore, the conservation of IGS regions in all strains, including the promoter elements of the *sxt* cluster that were identical at the nucleotide level, likely undergo similar transcriptional regulation. This suggests that the variation in PST production from toxic *D. circinale* strains has less to do with transcriptional regulation and more dependent on other regulatory mechanisms such as post-transcriptional. Underlining this is the ability of the four strains having the genes necessary to produce all the PSTs mentioned (Table 3) yet producing unique toxin profiles.

A previous study by Cullen et al. (2018) demonstrated that PST profiles in *S. crispum* strains were correlated to the presence of functional *sxt* tailoring genes, while our results suggest that PST profiles are also regulated by factors other than the *sxt* gene cluster. An investigation into the effects of environmental variables (such as pH, light and nutrient availability) on toxic *D. circinale* strains via transcriptomic and proteomic analyses may elucidate these regulatory factors. Moreover, the functional characterisation of tailoring genes as well as other uncharacterised *sxt* genes is necessary for the confirmation of strain-specific PST biosynthesis pathways and would also facilitate the monitoring and management strategies of cHABs.

## Acknowledgements

The work was funded by the Australian Research Council. The authors would like to thank Cathy Johnston from the ANACC for providing cyanobacterial isolates used in this project. The authors declare no conflicts of interest in relation to this manuscript.

## Notes

### Competing Interest Statement

The authors have declared no competing interest.

## References

Alexova, R., Haynes, P. A., Ferrari, B. C. & Neilan, B. A. 2011. Comparative protein expression in different strains of the bloom-forming cyanobacterium *Microcystis aeruginosa*. Mol Cell Proteomics, 10, M110 003749.

Ballot, A., Cerasino, L., Hostyeva, V. & Cires, S. 2016. Variability in the *sxt* gene clusters of PSP toxin producing *Aphanizomenon gracile* strains from Norway, Spain, Germany and North America. PLoS One, 11, e0167552.

Bankevich, A., Nurk, S., Antipov, D., Gurevich, A. A., Dvorkin, M., Kulikov, A. S., Lesin, V. M., Nikolenko, S. I., Pham, S., Prjibelski, A. D., Pyshkin, A. V., Sirotkin, A. V., Vyahhi, N., Tesler, G., Alekseyev, M. A. & Pevzner, P. A. 2012. SPAdes: a new genome assembly algorithm and its applications to single-cell sequencing. J Comput Biol, 19, 455–77.

Bateman, A., Martin, M. J., Orchard, S., Magrane, M., Alpi, E., Bely, B., Bingley, M., Britto, R., Bursteinas, B., Busiello, G., Bye-A-Jee, H., Da Silva, A., DE Giorgi, M., Dogan, T., Castro, L. G., Garmiri, P., Georghiou, G., Gonzales, D., Gonzales, L., Hatton-Ellis, E., Ignatchenko, A., Ishtiaq, R., Jokinen, P., Joshi, V., Jyothi, D., Lopez, R., Luo, J., Lussi, Y., Macdougall, A., Madeira, F., Mahmoudy, M., Menchi, M., Nightingale, A., Onwubiko, J., Palka, B., Pichler, K., Pundir, S., Qi, G. Y., Raj, S., Renaux, A., Lopez, M. R., Saidi, R., Sawford, T., Shypitsyna, A., Speretta, E., Turner, E., Tyagi, N., Vasudev, P., Volynkin, V., Wardell, T., Warner, K., Watkins, X., Zaru, R., Zellner, H., Bridge, A., Xenarios, I., Poux, S., Redaschi, N., Aimo, L., Argoud-Puy, G., Auchincloss, A., Axelsen, K., Bansal, P., Baratin, D., Blatter, M. C., Bolleman, J., Boutet, E., Breuza, L., Casals-Casas, C., DE Castro, E., Coudert, E., Cuche, B., Doche, M., Dornevil, D., Estreicher, A., Famiglietti, L., Feuermann, M., Gasteiger, E., Gehant, S., Gerritsen, V., Gos, A., Gruaz, N., Hinz, U., Hulo, C., Hyka-Nouspikel, N., Jungo, F., Keller, G., Kerhornou, A., Lara, V., Lemercier, P., Lieberherr, D., Lombardot, T., Martin, X., Masson, P., Morgat, A., Neto, T. B., Paesano, S., Pedruzzi, I., Pilbout, S., Pozzato, M., et al. 2019. UniProt: a worldwide hub of protein knowledge. Nucleic Acids Res, 47, D506–D515.

Blin, K., Shaw, S., Steinke, K., Villebro, R., Ziemert, N., Lee, S. Y., Medema, M. H. & Weber, T. 2019. antiSMASH 5.0: updates to the secondary metabolite genome mining pipeline. Nucleic Acids Res, 47, W81–W87.

Bolch, C. J. S. & Blackburn, S. I. 1996. Isolation and purification of Australian isolates of the toxic cyanobacterium *Microcystis aeruginosa* Kutz. J Appl Phycol, 8, 5–13.

Bolger, A. M., Lohse, M. & Usadel, B. 2014. Trimmomatic: a flexible trimmer for Illumina sequence data. Bioinformatics, 30, 2114–20.

Bowling, L. 1994. Occurrence and possible causes of a severe cyanobacterial bloom in Lake Cargelligo, New-South-Wales. Aus J Mar Fresh Res, 45, 737–745.

Bowling, L. C. & Baker, P. D. 1996. Major cyanobacterial bloom in the Barwon-Darling river, Australia, in 1991, and underlying limnological conditions. Mar Fresh Res, 47, 643-657.

Carmichael, W. W., Evans, W. R., Yin, Q. Q., Bell, P. & Moczydlowski, E. 1997. Evidence for paralytic shellfish poisons in the freshwater cyanobacterium *Lyngbya wollei* (Farlow ex Gomont) comb. nov. Appl Environ Microbiol, 63, 3104–3110.

Caspi, R., Billington, R., Fulcher, C. A., Keseler, I. M., Kothari, A., Krummenacker, M., Latendresse, M., Midford, P. E., Ong, Q., Ong, W. K., Paley, S., Subhraveti, P. & Karp, P. D. 2018. The MetaCyc database of metabolic pathways and enzymes. Nucleic Acids Research, 46, D633–D639.

Chen, I. A., Chu, K., Palaniappan, K., Pillay, M., Ratner, A., Huang, J., Huntemann, M., Varghese, N., White, J. R., Seshadri, R., Smirnova, T., Kirton, E., Jungbluth, S. P., Woyke, T., Eloe-Fadrosh, E. A., Ivanova, N. N. & Kyrpides, N. C. 2019. IMG/M v.5.0: an integrated data management and comparative analysis system for microbial genomes and microbiomes. Nucleic Acids Res, 47, D666-D677.

Cullen, A., D’agostino, P. M., Mazmouz, R., Pickford, R., Wood, S. & Neilan, B. A. 2018. Insertions within the saxitoxin biosynthetic gene cluster result in differential toxin profiles. ACS Chem Biol, 13, 3107–3114.

D’agostino, P. M., Song, X., Neilan, B. A. & Moffitt, M. C. 2014. Comparative proteomics reveals that a saxitoxin-producing and a nontoxic strain of *Anabaena circinalis* are two different ecotypes. J Proteome Res, 13, 1474–84.

D’agostino, P. M., Song, X., Neilan, B. A. & Moffitt, M. C. 2016. Proteogenomics of a saxitoxin-producing and non-toxic strain of *Anabaena circinalis* (cyanobacteria) in response to extracellular NaCl and phosphate depletion. Environ Microbiol, 18, 461–76.

Eddy, S. R. 1998. Profile hidden Markov models. Bioinformatics, 14, 755–63.

El-Gebali, S., Mistry, J., Bateman, A., Eddy, S. R., Luciani, A., Potter, S. C., Qureshi, M., Richardson, L. J., Salazar, G. A., Smart, A., Sonnhammer, E. L. L., Hirsh, L., Paladin, L., Piovesan, D., Tosatto, S. C. E. & Finn, R. D. 2019. The Pfam protein families database in 2019. Nucleic Acids Res, 47, D427–D432.

Ginn, H. P., Pearson, L. A. & Neilan, B. A. 2010. NtcA from *Microcystis aeruginosa* PCC 7806 is autoregulatory and binds to the microcystin promoter. Appl Environ Microbiol, 76, 4362–4368.

Haft, D. H., Loftus, B. J., Richardson, D. L., Yang, F., Eisen, J. A., Paulsen, I. T. & White, O. 2001. TIGRFAMs: a protein family resource for the functional identification of proteins. Nucleic Acids Res, 29, 41–3.

Humpage, A. R., Rositano, J., Bretag, A. H., Brown, R., Baker, P. D., Nicholson, B. C. & Steffensen, D. A. 1994. Paralytic shellfish poisons from Australian cyanobacterial blooms. Aus J Mar Fresh Res, 45, 761–771.

Kanehisa, M. & Goto, S. 2000. KEGG: kyoto encyclopedia of genes and genomes. Nucleic Acids Res, 28, 27–30.

Kang, D. D., Froula, J., Egan, R. & Wang, Z. 2015. Metabat, an efficienttool for accurately reconstructing single genomes from complex microbial communities. Peerj, 3, e1165.

Katoh, K. & Standley, D. M. 2013. MAFFT multiple sequence alignment software version 7: improvements in performance and usability. Mol Biol Evol, 30, 772–80.

Kearse, M., Moir, R., Wilson, A., Stones-Havas, S., Cheung, M., Sturrock, S., Buxton, S., Cooper, A., Markowitz, S., Duran, C., Thierer, T., Ashton, B., Meintjes, P. & Drummond, A. 2012. Geneious Basic: an integrated and extendable desktop software platform for the organization and analysis of sequence data. Bioinformatics, 28, 1647–9.

Kellmann, R., Mihali, T. K., Jeon, Y. J., Pickford, R., Pomati, F. & Neilan, B. A. 2008a. Biosynthetic intermediate analysis and functional homology reveal a saxitoxin gene cluster in cyanobacteria. Appl Environ Microbiol, 74, 4044–53.

Kellmann, R., Mihali, T. K. & Neilan, B. A. 2008b. Identification of a saxitoxin biosynthesis gene with a history of frequent horizontal gene transfers. J Mol Evol, 67, 526–38.

Kumar, S., Stecher, G., Li, M., Knyaz, C. & Tamura, K. 2018. MEGA X: Molecular evolutionary genetics analysis across computing platforms. Mol Biol and Evol, 35, 1547–1549.

Kurtz, S., Phillippy, A., Delcher, A. L., Smoot, M., Shumway, M., Antonescu, C. & Salzberg, S. L. 2004. Versatile and open software for comparing large genomes. Genome Biol, 5, R12.

Langmead, B. & Salzberg, S. L. 2012. Fast gapped-read alignment with Bowtie 2. Nat Methods, 9, 357–9.

Li, H., Handsaker, B., Wysoker, A., Fennell, T., Ruan, J., Homer, N., Marth, G., Abecasis, G., Durbin, R. & GENOME PROJECT DATA Processing, S. 2009. The Sequence alignment/map format and SAMtools. Bioinformatics, 25, 2078-9.

Li, X., Dreher, T. W. & Li, R. 2016. An overview of diversity, occurrence, genetics and toxin production of bloom-forming *Dolichospermum* (Anabaena) species. Harmful Algae, 54, 54–68.

Llewellyn, L. E., Negri, A. P., Doyle, J., Baker, P. D., Beltran, E. C. & Neilan, B. A. 2001. Radioreceptor assays for sensitive detection and quantitation of saxitoxin and its analogues from strains of the freshwater cyanobacterium, *Anabaena circinalis*. Environmental Science & Technology, 35, 1445–1451.

Lukowski, A. L., Ellinwood, D. C., Hinze, M. E., Deluca, R. J., Du Bois, J., Hall, S. & Narayan, A. R. H. 2018. C-H Hydroxylation in paralytic shellfish toxin biosynthesis. J Am Chem Soc, 140, 11863–11869.

Martin-Luna, B., Sevilla, E., Hernandez, J. A., Bes, M. T., Fillat, M. F. & Peleato, M. L. 2006. Fur from *Microcystis aeruginosa* binds *in vitro* promoter regions of the microcystin biosynthesis gene cluster. Phytochemistry, 67, 876–881.

Mihali, T. K., Carmichael, W. W. & Neilan, B. A. 2011. A putative gene cluster from a *Lyngbya wollei* bloom that encodes paralytic shellfish toxin biosynthesis. PLoS One, 6, e14657.

Mihali, T. K., Kellmann, R. & Neilan, B. A. 2009. Characterisation of the paralytic shellfish toxin biosynthesis gene clusters in *Anabaena circinalis* AWQC131C and *Aphanizomenon* sp. NH-5. BMC Biochem, 10, 8.

Morin, N., Vallaeys, T., Hendrickx, L., Natalie, L. & Wilmotte, A. 2010. An efficient DNA isolation protocol for filamentous cyanobacteria of the genus *Arthrospira*. J Microbiol Methods, 80, 148–54.

Okonechnikov, K., Golosova, O., Fursov, M. & Team, U. 2012. UniPro UGENE: a unified bioinformatics toolkit. Bioinformatics, 28, 1166–7.

Ongley, S. E., Pengelly, J. J. & Neilan, B. A. 2016. Elevated Na(+) and pH influence the production and transport of saxitoxin in the cyanobacteria *Anabaena circinalis* AWQC131C and *Cylindrospermopsis raciborskii* T3. Environ Microbiol, 18, 427–38.

Paerl, H. W. & Otten, T. G. 2013. Harmful cyanobacterial blooms: causes, consequences, and controls. Microb Ecol, 65, 995–1010.

Parks, D. H., Imelfort, M., Skennerton, C. T., Hugenholtz, P. & Tyson, G. W. 2015. CheckM: assessing the quality of microbial genomes recovered from isolates, single cells, and metagenomes. Genome Res, 25, 1043–55.

Pearson, L. A., Dittmann, E., Mazmouz, R., Ongley, S. E., D’agostino, P. M. & Neilan, B. A. 2016. The genetics, biosynthesis and regulation of toxic specialized metabolites of cyanobacteria. Harmful Algae, 54, 98–111.

Pereyra, J. P. A., D’agostino, P. M., Mazmouz, R., Woodhouse, J. N., Pickford, R., Jameson, I. & Neilan, B. A. 2017. Molecular and morphological survey of saxitoxin-producing cyanobacterium *Dolichospermum circinale* (*Anabaena circinalis*) isolated from geographically distinct regions of Australia. Toxicon, 138, 68–77.

Rodriguez, R. L., Gunturu, S., Harvey, W. T., Rossello-Mora, R., Tiedje, J. M., Cole, J. R. & Konstantinidis, K. T. 2018. The Microbial Genomes Atlas (MiGA) webserver: taxonomic and gene diversity analysis of Archaea and Bacteria at the whole genome level. Nucleic Acids Res, 46, W282–W288.

Sinha, R., Pearson, L. A., Davis, T. W., Muenchhoff, J., Pratama, R., Jex, A., Burford, M. A. & Neilan, B. A. 2014. Comparative genomics of *Cylindrospermopsis raciborskii* strains with differential toxicities. BMC Genomics, 15, 83.

Smith, F. M., Wood, S. A., Van Ginkel, R., Broady, P. A. & Gaw, S. 2011. First report of saxitoxin production by a species of the freshwater benthic cyanobacterium, *Scytonema agardhi*. Toxicon, 57, 566–73.

Soto-Liebe, K., Lopez-Cortes, X. A., Fuentes-Valdes, J. J., Stucken, K., Gonzalez-Nilo, F. & Vasquez, M. 2013. *In silico* analysis of putative paralytic shellfish poisoning toxins export proteins in Cyanobacteria. PloS One, 8.

Soto-Liebe, K., Murillo, A. A., Krock, B., Stucken, K., Fuentes-Valdes, J. J., Trefault, N., Cembella, A. & Vasquez, M. 2010. Reassessment of the toxin profile of *Cylindrospermopsis raciborskii* T3 and function of putative sulfotransferases in synthesis of sulfated and sulfonated PSP toxins. Toxicon, 56, 1350–61.

Stucken, K., John, U., Cembella, A., Murillo, A. A., Soto-Liebe, K., Fuentes-Valdes, J. J., Friedel, M., Plominsky, A. M., Vasquez, M. & Glockner, G. 2010. The smallest known genomes of multicellular and toxic cyanobacteria: comparison, minimal gene sets for linked traits and the evolutionary implications. PLoS One, 5, e9235.

Tatusov, R. L., Galperin, M. Y., Natale, D. A. & Koonin, E. V. 2000. The COG database: a tool for genome-scale analysis of protein functions and evolution. Nucleic Acids Res, 28, 33–6.

Velzeboer, R. M. A., Baker, P. D. & Rositano, J. 2001. Saxitoxins associated with the growth of the cyanobacterium *Anabaena circinalis* (Nostocales, Cyanophyta) under varying sources and concentrations of nitrogen. Phycologia, 40, 305–312.

Velzeboer, R. M. A., Baker, P. D., Rositano, J., Heresztyn, T., Codd, G. A. & Raggett, S. L. 2000. Geographical patterns of occurrence and composition of saxitoxins in the cyanobacterial genus *Anabaena* (Nostocales, Cyanophyta) in Australia. Phycologia, 39, 395–407.

Wiese, M., D’agostino, P. M., Mihali, T. K., Moffitt, M. C. & Neilan, B. A. 2010. Neurotoxic alkaloids: saxitoxin and its analogs. Mar Drugs, 8, 2185–211.

Wilde, A. & Hihara, Y. 2016. Transcriptional and posttranscriptional regulation of cyanobacterial photosynthesis. Biochimica Et Biophysica Acta-Bioenergetics, 1857, 296–308.

Willis, A., Woodhouse, J. N., Ongley, S. E., Jex, A. R., Burford, M. A. & Neilan, B. A. 2018. Genome variation in nine co-occurring toxic *Cylindrospermopsis raciborskii* strains. Harmful Algae, 73, 157–166.

Yoon, S. H., Ha, S. M., Lim, J., Kwon, S. & Chun, J. 2017. A large-scale evaluation of algorithms to calculate average nucleotide identity. Antonie Van Leeuwenhoek, 110, 1281–1286.

Zhang, X. W., Zhao, F. Q., Guan, X. G., Yang, Y., Liang, C. W. & Qin, S. 2007. Genome-wide survey of putative serine/threonine protein kinases in cyanobacteria. BMC Genomics, 8.

