## Supplemental Data for "Comparative genomics of *Dolichospermum circinale* strains with differential paralytic shellfish toxin profiles"

### 1 Phylogenetic analysis

A specific set of 198 marker proteins were identified from all bacterial genomes in the NCBI database, from 7,315 genomes. These marker proteins were present in ≥95% of genomes in the database, and existed as single-copy genes in ≥95% of cases. Accession numbers are from the Pfam (PF) (El-Gebali et al., 2019) or TIGRFAM (TIGR) (Haft et al., 2001) databases.

**Supplementary Table S1**: 198 marker proteins used for phylogenetic analysis.

| Number | Name | Accession No. | Description |
|---|---|---|---|
| 1 | AAA_17 | PF13207.5 | AAA domain |
| 2 | ADK | PF00406.21 | Adenylate kinase |
| 3 | B5 | PF03484.14 | tRNA synthetase B5 domain |
| 4 | CoaE | PF1121.19 | Dephospho-CoA kinase |
| 5 | Competence | PF03772.15 | Competence protein |
| 6 | CTP_synth_N | PF06418.13 | CTP synthase N-terminus |
| 7 | DNA_pol3_beta | PF00712.18 | DNA polymerase III beta subunit, N-terminal domain |
| 8 | DNA_pol3_beta_3 | PF02768.14 | DNA polymerase III beta subunit, C-terminal domain |
| 9 | EF_TS | PF00889.18 | Elongation factor TS |
| 10 | FDX-ACB | PF03147.13 | Ferredoxin-fold anticodon binding domain |
| 11 | Flavokinase | PF01687.16 | Riboflavin kinase |
| 12 | FtsZ_C | PF12327.7 | FtsZ family, C-terminal domain |
| 13 | GidB | PF02527.14 | rRNA small subunit methyltransferase G |
| 14 | GrpE | PF01025.18 | GrpE |
| 15 | GTP1_OBG | PF01018.21 | GTP1/OBG |
| 16 | IF-2 | PF11987.7 | Translation-initiation factor 2 |
| 17 | IF3_C | PF00707.21 | Translation initiation factor IF-3, C-terminal domain |
| 18 | IPPT | PF01715.16 | IPP transferase |
| 19 | KH_5 | PF13184.5 | NusA-like KH domain |
| 20 | KH_dom-like | PF14714.5 | KH-domain-like of EngA bacterial GTPase enzymes, C-terminal |
| 21 | LepA_C | PF06421.11 | GTP-binding protein LepA C-terminus |
| 22 | Methyltransf_5 | PF01795.18 | MraW methylase family |
| 23 | MurB_C | PF02873.15 | UDP-N-acetylenolpyruvoylglucosamine reductase, C-terminal domain |
| 24 | Peptidase_M50 | PF02163.21 | Peptidase family M50 |
| 25 | Pept_tRNA_hydro | PF01195.18 | Peptidyl-tRNA hydrolase |

| Number | Name | Accession No. | Description |
| --- | --- | --- | --- |
| 26 | PGK | PF00162.18 | Phosphoglycerate kinase |
| 27 | Phe_tRNA-synt_N | PF02912.17 | Aminoacyl tRNA synthetase class II, N-terminal domain |
| 28 | RBFA | PF02033.17 | Ribosome-binding factor A |
| 29 | RecA | PF00154.20 | recA bacterial DNA recombination protein |
| 30 | Ribonucleas_3_3 | PF14622.5 | Ribonuclease-III-like |
| 31 | Ribosomal_L1 | PF00687.20 | Ribosomal protein L1p/L10e family |
| 32 | Ribosomal_L10 | PF00466.19 | Ribosomal protein L10 |
| 33 | Ribosomal_L11 | PF00298.18 | Ribosomal protein L11, RNA binding domain |
| 34 | Ribosomal_L11_N | PF03946.13 | Ribosomal protein L11, N-terminal domain |
| 35 | Ribosomal_L12 | PF00542.18 | Ribosomal protein L7/L12 C-terminal domain |
| 36 | Ribosomal_L12_N | PF16320.4 | Ribosomal protein L7/L12 dimerisation domain |
| 37 | Ribosomal_L13 | PF00572.17 | Ribosomal protein L13 |
| 38 | Ribosomal_L14 | PF00238.18 | Ribosomal protein L14p/L23e |
| 39 | Ribosomal_L16 | PF00252.17 | Ribosomal protein L16p/L10e |
| 40 | Ribosomal_L17 | PF01196.18 | Ribosomal protein L17 |
| 41 | Ribosomal_L18p | PF00861.21 | Ribosomal L18 of archaea, bacteria, mitoch. and chloroplast |
| 42 | Ribosomal_L19 | PF01245.19 | Ribosomal protein L19 |
| 43 | Ribosomal_L2 | PF00181.22 | Ribosomal Proteins L2, RNA binding domain |
| 44 | Ribosomal_L20 | PF00453.17 | Ribosomal protein L20 |
| 45 | Ribosomal_L21p | PF00829.20 | Ribosomal prokaryotic L21 protein |
| 46 | Ribosomal_L22 | PF00237.18 | Ribosomal protein L22p/L17e |
| 47 | Ribosomal_L23 | PF00276.19 | Ribosomal protein L23 |
| 48 | Ribosomal_L27 | PF01016.18 | Ribosomal L27 protein |
| 49 | Ribosomal_L27A | PF00828.18 | Ribosomal proteins 50S-L15, 50S-L18e, 60S-L27A |
| 50 | Ribosomal_L29 | PF00831.22 | Ribosomal L29 protein |
| 51 | Ribosomal_L2_C | PF03947.17 | Ribosomal Proteins L2, C-terminal domain |
| 52 | Ribosomal_L3 | PF00297.21 | Ribosomal protein L3 |

| Number | Name | Accession No. | Description |
|---|---|---|---|
| 53 | Ribosomal_L32p | PF01783.22 | Ribosomal L32p protein family |
| 54 | Ribosomal_L35p | PF01632.18 | Ribosomal protein L35 |
| 55 | Ribosomal_L4 | PF00573.21 | Ribosomal protein L4/L1 family |
| 56 | Ribosomal_L5 | PF00281.18 | Ribosomal protein L5 |
| 57 | Ribosomal_L5_C | PF00673.20 | ribosomal L5P family C-terminus |
| 58 | Ribosomal_L6 | PF00347.22 | Ribosomal protein L6 |
| 59 | Ribosomal_L9_C | PF03948.13 | Ribosomal protein L9, C-terminal domain |
| 60 | Ribosomal_L9_N | PF01281.18 | Ribosomal protein L9, N-terminal domain |
| 61 | Ribosomal_S10 | PF00338.21 | Ribosomal protein S10p/S20e |
| 62 | Ribosomal_S13 | PF00416.21 | Ribosomal protein S13/S18 |
| 63 | Ribosomal_S15 | PF00312.21 | Ribosomal protein S15 |
| 64 | Ribosomal_S16 | PF00886.18 | Ribosomal protein S16 |
| 65 | Ribosomal_S17 | PF00366.19 | Ribosomal protein S17 |
| 66 | Ribosomal_S19 | PF00203.20 | Ribosomal protein S19 |
| 67 | Ribosomal_S2 | PF00318.19 | Ribosomal protein S2 |
| 68 | Ribosomal_S3_C | PF00189.19 | Ribosomal protein S3, C-terminal domain |
| 69 | Ribosomal_S4 | PF00163.18 | Ribosomal protein S4/S9 N-terminal domain |
| 70 | Ribosomal_S5 | PF00333.19 | Ribosomal protein S5, N-terminal domain |
| 71 | Ribosomal_S5_C | PF03719.14 | Ribosomal protein S5, C-terminal domain |
| 72 | Ribosomal_S6 | PF01250.16 | Ribosomal protein S6 |
| 73 | Ribosomal_S7 | PF00177.20 | Ribosomal protein S7p/S5e |
| 74 | Ribosomal_S8 | PF00410.18 | Ribosomal protein S8 |
| 75 | Ribosomal_S9 | PF00380.18 | Ribosomal protein S9/S16 |
| 76 | Ribosom_S12_S23 | PF00164.24 | Ribosomal protein S12/S23 |
| 77 | RNA_pol_A_bac | PF01000.25 | RNA polymerase Rpb3/RpoA insert domain |
| 78 | RNA_pol_A_CTD | PF03118.14 | Bacterial RNA polymerase, alpha chain C terminal domain |
| 79 | RNA_pol_L | PF01193.23 | RNA polymerase Rpb3/Rpb11 dimerisation domain |

| Number | Name | Accession No. | Description |
|---|---|---|---|
| 80 | RNA_pol_Rpb1_1 | PF04997.11 | RNA polymerase Rpb1, domain 1 |
| 81 | RNA_pol_Rpb1_2 | PF00623.19 | RNA polymerase Rpb1, domain 2 |
| 82 | RNA_pol_Rpb1_3 | PF04983.17 | RNA polymerase Rpb1, domain 3 |
| 83 | RNA_pol_Rpb1_5 | PF04998.16 | RNA polymerase Rpb1, domain 5 |
| 84 | RNA_pol_Rpb2_1 | PF04563.14 | RNA polymerase beta subunit |
| 85 | RNA_pol_Rpb2_3 | PF04565.15 | RNA polymerase Rpb2, domain 3 |
| 86 | RNA_pol_Rpb2_45 | PF10385.8 | RNA polymerase beta subunit external 1 domain |
| 87 | RNA_pol_Rpb2_6 | PF00562.27 | RNA polymerase Rpb2, domain 6 |
| 88 | RNA_pol_Rpb2_7 | PF04560.19 | RNA polymerase Rpb2, domain 7 |
| 89 | RRF | PF01765.18 | Ribosome recycling factor |
| 90 | RuvB_C | PF05491.12 | Holliday junction DNA helicase ruvB C-terminus |
| 91 | RuvX | PF03652.14 | Holliday junction resolvase |
| 92 | S-AdoMet_synt_C | PF02773.15 | S-adenosylmethionine synthetase, C-terminal domain |
| 93 | S-AdoMet_synt_M | PF02772.15 | S-adenosylmethionine synthetase, central domain |
| 94 | S-AdoMet_synt_N | PF00438.19 | S-adenosylmethionine synthetase, N-terminal domain |
| 95 | Seryl_tRNA_N | PF02403.21 | Seryl-tRNA synthetase N-terminal domain |
| 96 | SmpB | PF01668.17 | SmpB protein |
| 97 | SRP_SPB | PF02978.18 | Signal peptide binding domain |
| 98 | Toprim_N | PF08275.10 | DNA primase catalytic core, N-terminal domain |
| 99 | TRCF | PF03461.14 | TRCF domain |
| 100 | Trigger_N | PF05697.12 | Bacterial trigger factor protein (TF) |
| 101 | tRNA-synt_1d | PF00750.18 | tRNA synthetases class I (R) |
| 102 | tRNA-synt_1_2 | PF13603.5 | Leucyl-tRNA synthetase, Domain 2 |
| 103 | tRNA-synt_2d | PF01409.19 | tRNA synthetases class II core domain (F) |
| 104 | tRNA_m1G_MT | PF01746.20 | tRNA (Guanine-1)-methyltransferase |
| 105 | tRNA_Me_trans | PF03054.15 | tRNA methyl transferase |
| 106 | TruB_N | PF01509.17 | TruB family pseudouridylate synthase (N terminal domain) |

| Number | Name | Accession No. | Description |
| --- | --- | --- | --- |
| 107 | TsaE | PF02367.16 | Threonylcarbamoyl adenosine biosynthesis protein TsaE |
| 108 | Tubulin | PF00091.24 | Tubulin/FtsZ family, GTPase domain |
| 109 | UPF0054 | PF02130.16 | Uncharacterized protein family UPF0054 |
| 110 | UvrB | PF12344.7 | Ultra-violet resistance protein B |
| 111 | UvrC_HhH_N | PF08459.10 | UvrC Helix-hairpin-helix N-terminal |
| 112 | YchF-GTPase_C | PF06071.12 | Protein of unknown function (DUF933) |
| 113 | zf-CHC2 | PF01807.19 | CHC2 zinc finger |
| 114 | TIGR00001 | TIGR00001 | rpmI_bact: ribosomal protein bL35 |
| 115 | TIGR00002 | TIGR00002 | S16: ribosomal protein bS16 |
| 116 | TIGR00006 | TIGR00006 | TIGR00006: 16S rRNA (cytosine(1402)-N(4))-methyltransferase |
| 117 | TIGR00012 | TIGR00012 | L29: ribosomal protein uL29 |
| 118 | TIGR00043 | TIGR00043 | TIGR00043: rRNA maturation RNase YbeY |
| 119 | TIGR00059 | TIGR00059 | L17: ribosomal protein bL17 |
| 120 | TIGR00060 | TIGR00060 | L18_bact: ribosomal protein uL18 |
| 121 | TIGR00061 | TIGR00061 | L21: ribosomal protein bL21 |
| 122 | TIGR00062 | TIGR00062 | L27: ribosomal protein bL27 |
| 123 | TIGR00065 | TIGR00065 | ftsZ: cell division protein FtsZ |
| 124 | TIGR00082 | TIGR00082 | rbfA: ribosome-binding factor A |
| 125 | TIGR00084 | TIGR00084 | ruvA: Holliday junction DNA helicase RuvA |
| 126 | TIGR00086 | TIGR00086 | smpB: SsrA-binding protein |
| 127 | TIGR00088 | TIGR00088 | trmD: tRNA (guanine(37)-N(1))-methyltransferase |
| 128 | TIGR00115 | TIGR00115 | tig: trigger factor |
| 129 | TIGR00116 | TIGR00116 | tsf: translation elongation factor Ts |
| 130 | TIGR00121 | TIGR00121 | birA_ligase: biotin--[acetyl-CoA-carboxylase] ligase |
| 131 | TIGR00138 | TIGR00138 | rsmG_gidB: 16S rRNA (guanine(527)-N(7))-methyltransferase RsmG |
| 132 | TIGR00150 | TIGR00150 | T6A_YjeE: tRNA threonylcarbamoyl adenosine modification protein YjeE |
| 133 | TIGR00152 | TIGR00152 | TIGR00152: dephospho-CoA kinase |

| Number | Name | Accession No. | Description |
|---|---|---|---|
| 134 | TIGR00158 | TIGR00158 | L9: ribosomal protein bL9 |
| 135 | TIGR00166 | TIGR00166 | S6: ribosomal protein bS6 |
| 136 | TIGR00168 | TIGR00168 | infC: translation initiation factor IF-3 |
| 137 | TIGR00174 | TIGR00174 | miaA: tRNA dimethylallyltransferase |
| 138 | TIGR00179 | TIGR00179 | murB: UDP-N-acetylenolpyruvoylglucosamine reductase |
| 139 | TIGR00250 | TIGR00250 | RNAse_H_YqgF: putative transcription antitermination factor YqgF |
| 140 | TIGR00420 | TIGR00420 | trmU: tRNA (5-methylaminomethyl-2-thiouridylate)-methyltransferase |
| 141 | TIGR00425 | TIGR00425 | CBF5: putative rRNA pseudouridine synthase |
| 142 | TIGR00431 | TIGR00431 | TruB: tRNA pseudouridine(55) synthase |
| 143 | TIGR00447 | TIGR00447 | pth: aminoacyl-tRNA hydrolase |
| 144 | TIGR00468 | TIGR00468 | pheS: phenylalanine--tRNA ligase, alpha subunit |
| 145 | TIGR00469 | TIGR00469 | pheS_mito: phenylalanine--tRNA ligase |
| 146 | TIGR00471 | TIGR00471 | pheT_arch: phenylalanine--tRNA ligase, beta subunit |
| 147 | TIGR00496 | TIGR00496 | frr: ribosome recycling factor |
| 148 | TIGR00615 | TIGR00615 | recR: recombination protein RecR |
| 149 | TIGR00855 | TIGR00855 | L12: ribosomal protein bL12 |
| 150 | TIGR00952 | TIGR00952 | S15_bact: ribosomal protein uS15 |
| 151 | TIGR00981 | TIGR00981 | rpsL_bact: ribosomal protein uS12 |
| 152 | TIGR01008 | TIGR01008 | uS3_euk_arch: ribosomal protein uS3 |
| 153 | TIGR01009 | TIGR01009 | rpsC_bact: ribosomal protein uS3 |
| 154 | TIGR01011 | TIGR01011 | rpsB_bact: ribosomal protein uS2 |
| 155 | TIGR01017 | TIGR01017 | rpsD_bact: ribosomal protein uS4 |
| 156 | TIGR01020 | TIGR01020 | uS5_euk_arch: ribosomal protein uS5 |
| 157 | TIGR01021 | TIGR01021 | rpsE_bact: ribosomal protein uS5 |
| 158 | TIGR01024 | TIGR01024 | rplS_bact: ribosomal protein bL19 |
| 159 | TIGR01025 | TIGR01025 | uS19_arch: ribosomal protein uS19 |
| 160 | TIGR01028 | TIGR01028 | uS7_euk_arch: ribosomal protein uS7 |

| Number | Name | Accession No. | Description |
|---|---|---|---|
| 161 | TIGR01029 | TIGR01029 | rpsG_bact: ribosomal protein uS7 |
| 162 | TIGR01032 | TIGR01032 | rplT_bact: ribosomal protein bL20 |
| 163 | TIGR01044 | TIGR01044 | rplV_bact: ribosomal protein uL22 |
| 164 | TIGR01046 | TIGR01046 | uS10_euk_arch: ribosomal protein uS10 |
| 165 | TIGR01049 | TIGR01049 | rpsJ_bact: ribosomal protein uS10 |
| 166 | TIGR01050 | TIGR01050 | rpsS_bact: ribosomal protein uS19 |
| 167 | TIGR01066 | TIGR01066 | rplM_bact: ribosomal protein uL13 |
| 168 | TIGR01067 | TIGR01067 | rplN_bact: ribosomal protein uL14 |
| 169 | TIGR01071 | TIGR01071 | rplO_bact: ribosomal protein uL15 |
| 170 | TIGR01079 | TIGR01079 | rplX_bact: ribosomal protein uL24 |
| 171 | TIGR01164 | TIGR01164 | rplP_bact: ribosomal protein uL16 |
| 172 | TIGR01169 | TIGR01169 | rplA_bact: ribosomal protein uL1 |
| 173 | TIGR01170 | TIGR01170 | rplA_mito: ribosomal protein uL1, mitochondrial |
| 174 | TIGR01171 | TIGR01171 | rplB_bact: ribosomal protein uL2 |
| 175 | TIGR01351 | TIGR01351 | adk: adenylate kinase |
| 176 | TIGR01359 | TIGR01359 | UMP_CMP_kin_fam: UMP-CMP kinase family |
| 177 | TIGR01360 | TIGR01360 | aden_kin_iso1: adenylate kinase |
| 178 | TIGR01632 | TIGR01632 | L11_bact: ribosomal protein uL11 |
| 179 | TIGR01953 | TIGR01953 | NusA: transcription termination factor NusA |
| 180 | TIGR02012 | TIGR02012 | tigrfam_recA: protein RecA |
| 181 | TIGR02013 | TIGR02013 | rpoB: DNA-directed RNA polymerase, beta subunit |
| 182 | TIGR02027 | TIGR02027 | rpoA: DNA-directed RNA polymerase, alpha subunit |
| 183 | TIGR02191 | TIGR02191 | RNaseIII: ribonuclease III |
| 184 | TIGR02386 | TIGR02386 | rpoC_TIGR: DNA-directed RNA polymerase, beta' subunit |
| 185 | TIGR02387 | TIGR02387 | rpoC1_cyan: DNA-directed RNA polymerase, gamma subunit |
| 186 | TIGR02388 | TIGR02388 | rpoC2_cyan: DNA-directed RNA polymerase, beta'' subunit |
| 187 | TIGR02389 | TIGR02389 | RNA_pol_rpoA2: DNA-directed RNA polymerase, subunit A'' |

| Number | Name | Accession No. | Description |
|---|---|---|---|
| 188 | TIGR02390 | TIGR02390 | RNA_pol_rpoA1: DNA-directed RNA polymerase subunit A' |
| 189 | TIGR03625 | TIGR03625 | L3_bact: 50S ribosomal protein uL3 |
| 190 | TIGR03627 | TIGR03627 | uS9_arch: ribosomal protein uS9 |
| 191 | TIGR03631 | TIGR03631 | uS13_bact: ribosomal protein uS13 |
| 192 | TIGR03635 | TIGR03635 | uS17_bact: ribosomal protein uS17 |
| 193 | TIGR03653 | TIGR03653 | uL6_arch: ribosomal protein uL6 |
| 194 | TIGR03654 | TIGR03654 | L6_bact: ribosomal protein uL6 |
| 195 | TIGR03670 | TIGR03670 | rpoB_arch: DNA-directed RNA polymerase subunit B |
| 196 | TIGR03673 | TIGR03673 | uL14_arch: 50S ribosomal protein uL14 |
| 197 | TIGR03722 | TIGR03722 | arch_KAE1: universal archaeal protein Kae1 |
| 198 | TIGR03953 | TIGR03953 | rplD_bact: 50S ribosomal protein uL4 |

#### 2 Taxonomic analysis

The genomes of the four strains in this study (ACBU01, ACMB03, ACMB13 and FSS-124) were compared at the nucleotide level via average nucleotide identity (ANI; Figure S1), and at the amino acid level via average amino acid identity (AAI; Figure S2), with the genomes of other strains as described in the Materials and Methods. All numbers in the heatmap figures are listed as percentages, and the four genomes from this study are labelled in bold. The species similarity cutoff for ANI and the taxonomic identity standards for AAI were as described by Konstantinidis et al. (2017) and were used to colour code the figures, with the remainder of the ANI value cutoffs implemented manually for ease of comparison.

| | *Agrobacterium fabrum* str. C58 | *Anabaena* sp. 90 chromosome chANA01 | *Anabaena* sp. 90 chromosome chANA02 | *Anabaena* sp. WA102 | *Anabaena variabilis* ATCC 29413 | *Anabaenopsis circularis* NIES 21 | *Aulosira laxa* NIES 50 | *Cylindrospermum* sp. NIES 4074 | ***Dolichospermum circinale* ACBU01** | ***Dolichospermum circinale* ACMB03** | ***Dolichospermum circinale* ACMB13** | *Dolichospermum circinale* AWQC131C | *Dolichospermum circinale* AWQC310F | ***Dolichospermum circinale* FSS-124** | *Dolichospermum compactum* NIES 806 | *Escherichia coli* str. K12 substr. MG1655 | *Gloeobacter violaceus* PCC 7421 | *Nostoc azollae* 0708 | *Nostoc carneum* NIES 2107 | *Nostoc sphaeroides* strain Kutzing | *Scytonema crispum* CAWBG524 |
|---|---|---|---|---|---|---|---|---|---|---|---|---|---|---|---|---|---|---|---|---|---|
| *Anabaena* sp. 90 chromosome chANA01 | 70.36 | | | | | | | | | | | | | | | | | | | | |
| *Anabaena* sp. 90 chromosome chANA02 | 61.80 | 81.22 | | | | | | | | | | | | | | | | | | | |
| *Anabaena* sp. WA102 | 69.05 | 90.66 | 90.13 | | | | | | | | | | | | | | | | | | |
| *Anabaena variabilis* ATCC 29413 | 66.82 | 74.54 | 74.70 | 74.73 | | | | | | | | | | | | | | | | | |
| *Anabaenopsis circularis* NIES 21 | 68.58 | 74.86 | 74.81 | 74.88 | 76.87 | | | | | | | | | | | | | | | | |
| *Aulosira laxa* NIES 50 | 66.73 | 74.77 | 75.10 | 74.81 | 76.10 | 76.16 | | | | | | | | | | | | | | | |
| *Cylindrospermum* sp. NIES 4074 | 67.97 | 76.56 | 76.65 | 76.53 | 76.11 | 76.56 | 76.87 | | | | | | | | | | | | | | |
| ***Dolichospermum circinale* ACBU01** | 69.08 | 85.40 | 84.93 | 85.12 | 74.42 | 74.52 | 74.58 | 76.08 | | | | | | | | | | | | | |
| ***Dolichospermum circinale* ACMB03** | 65.43 | 85.38 | 84.89 | 85.21 | 74.27 | 74.55 | 74.67 | 76.00 | 99.92 | | | | | | | | | | | | |
| ***Dolichospermum circinale* ACMB13** | 62.28 | 85.38 | 84.94 | 85.07 | 74.36 | 74.61 | 74.74 | 75.96 | 99.91 | 99.93 | | | | | | | | | | | |
| *Dolichospermum circinale* AWQC131C | 69.51 | 85.49 | 85.19 | 85.12 | 74.36 | 74.41 | 74.58 | 75.99 | 99.83 | 99.84 | 99.83 | | | | | | | | | | |
| *Dolichospermum circinale* AWQC310F | 65.00 | 85.55 | 84.93 | 85.19 | 74.25 | 74.54 | 74.61 | 76.08 | 96.07 | 96.17 | 96.09 | 96.13 | | | | | | | | | |
| ***Dolichospermum circinale* FSS-124** | 63.07 | 85.46 | 85.00 | 85.12 | 74.31 | 74.46 | 74.48 | 76.00 | 98.41 | 98.40 | 98.42 | 98.43 | 96.04 | | | | | | | | |
| *Dolichospermum compactum* NIES 806 | 70.47 | 89.47 | 89.45 | 93.23 | 74.60 | 74.78 | 74.94 | 76.28 | 85.09 | 85.05 | 84.98 | 85.10 | 84.94 | 85.12 | | | | | | | |
| *Escherichia coli* str. K12 substr. MG1655 | 65.71 | 65.06 | 62.41 | 63.73 | 64.22 | 63.33 | 64.02 | 63.87 | 62.35 | 62.48 | 62.05 | 63.35 | 63.67 | 61.95 | 64.15 | | | | | | |
| *Gloeobacter violaceus* PCC 7421 | 65.93 | 64.45 | 64.62 | 64.54 | 64.76 | 64.70 | 64.14 | 65.32 | 64.43 | 64.26 | 64.44 | 64.51 | 64.78 | 64.16 | 64.43 | 63.47 | | | | | |
| *Nostoc azollae* 0708 | 72.07 | 78.40 | 77.97 | 78.66 | 74.78 | 75.25 | 75.26 | 77.00 | 77.76 | 77.90 | 78.01 | 77.88 | 77.89 | 77.99 | 78.35 | 64.86 | 65.22 | | | | |
| *Nostoc carneum* NIES 2107 | 65.24 | 74.76 | 75.24 | 74.65 | 76.17 | 76.21 | 93.84 | 77.02 | 74.52 | 74.46 | 74.51 | 74.59 | 74.59 | 74.45 | 74.81 | 63.34 | 64.84 | 75.13 | | | |
| *Nostoc sphaeroides* strain Kutzing | 67.90 | 75.41 | 75.57 | 75.32 | 76.90 | 77.12 | 77.63 | 79.39 | 74.87 | 74.97 | 74.99 | 74.90 | 74.96 | 74.85 | 75.37 | 63.45 | 64.89 | 75.90 | 77.50 | | |
| *Scytonema crispum* CAWBG524 | 62.86 | 74.25 | 74.44 | 74.15 | 75.14 | 75.68 | 76.38 | 77.13 | 74.28 | 74.23 | 74.29 | 74.26 | 74.09 | 74.19 | 74.33 | 63.24 | 64.88 | 74.93 | 76.30 | 78.00 | |
| *Scytonema crispum* CAWBG72 | 62.13 | 74.33 | 75.11 | 74.28 | 75.18 | 75.69 | 76.11 | 77.09 | 74.26 | 74.23 | 74.20 | 74.26 | 73.99 | 74.11 | 74.34 | 63.38 | 64.95 | 74.86 | 76.15 | 77.99 | 99.02 |

**ANI Values**

- >95%; Species
- 80% - 95%
- 65% - 80%
- 50% - 65%

**Figure S1**: Triangle heatmap of pairwise ANI calculations. All numbers in the heatmap are listed as percentages. The four genomes from this study are labelled in bold. The >95% cutoff for species similarity based on ANI is as described by Konstantinidis et al. (2017).

| | *Agrobacterium fabrum* str. C58 | *Anabaena* sp. 90 chromosome chANA01 | *Anabaena* sp. 90 chromosome chANA02 | *Anabaena* sp. WA102 | *Anabaena variabilis* ATCC 29413 | *Anabaenopsis circularis* NIES 21 | *Aulosira laxa* NIES 50 | *Cylindrospermum* sp. NIES 4074 | ***Dolichospermum circinale* ACBU01** | ***Dolichospermum circinale* ACMB03** | ***Dolichospermum circinale* ACMB13** | *Dolichospermum circinale* AWQC131C | *Dolichospermum circinale* AWQC310F | ***Dolichospermum circinale* FSS-124** | *Dolichospermum compactum* NIES 806 | *Escherichia coli* str. K12 substr. MG1655 | *Gloeobacter violaceus* PCC 7421 | *Nostoc azollae* 0708 | *Nostoc carneum* NIES 2107 | *Nostoc sphaeroides* strain Kutzing | *Scytonema crispum* CAWBG524 |
|---|---|---|---|---|---|---|---|---|---|---|---|---|---|---|---|---|---|---|---|---|---|
| *Anabaena* sp. 90 chromosome chANA01 | 42.08 | | | | | | | | | | | | | | | | | | | | |
| *Anabaena* sp. 90 chromosome chANA02 | 40.39 | 54.37 | | | | | | | | | | | | | | | | | | | |
| *Anabaena* sp. WA102 | 42.16 | 90.64 | 89.89 | | | | | | | | | | | | | | | | | | |
| *Anabaena variabilis* ATCC 29413 | 42 | 73.98 | 73.71 | 74.05 | | | | | | | | | | | | | | | | | |
| *Anabaenopsis circularis* NIES 21 | 42.49 | 73.61 | 74.02 | 73.89 | 78.07 | | | | | | | | | | | | | | | | |
| *Aulosira laxa* NIES 50 | 43.21 | 73.5 | 73.48 | 73.56 | 76.82 | 76.67 | | | | | | | | | | | | | | | |
| *Cylindrospermum* sp. NIES 4074 | 41.81 | 75.98 | 75.21 | 76.14 | 76.9 | 76.85 | 76.88 | | | | | | | | | | | | | | |
| ***Dolichospermum circinale* ACBU01** | 41.64 | 85.4 | 83.95 | 85.4 | 74.26 | 74.03 | 73.84 | 76.11 | | | | | | | | | | | | | |
| ***Dolichospermum circinale* ACMB03** | 41.64 | 85.39 | 83.92 | 85.39 | 74.24 | 74.03 | 73.85 | 76.1 | 100 | | | | | | | | | | | | |
| ***Dolichospermum circinale* ACMB13** | 41.63 | 85.4 | 84.06 | 85.4 | 74.25 | 74.06 | 73.86 | 76.18 | 99.99 | 100 | | | | | | | | | | | |
| *Dolichospermum circinale* AWQC131C | 41.65 | 85.38 | 84.11 | 85.43 | 74.28 | 74.12 | 73.91 | 76.16 | 99.99 | 99.98 | 99.98 | | | | | | | | | | |
| *Dolichospermum circinale* AWQC310F | 41.94 | 85.08 | 83.69 | 85.38 | 74.16 | 74.18 | 74.06 | 76.26 | 96.15 | 96.18 | 96.16 | 96.2 | | | | | | | | | |
| ***Dolichospermum circinale* FSS-124** | 41.66 | 85.08 | 84.38 | 85.28 | 74.18 | 74.04 | 73.92 | 76.11 | 98.33 | 98.34 | 98.34 | 98.33 | 96.11 | | | | | | | | |
| *Dolichospermum compactum* NIES 806 | 42.43 | 90.08 | 89.55 | 93.97 | 74.45 | 74.38 | 73.96 | 76.53 | 85.71 | 85.71 | 85.76 | 85.74 | 85.55 | 85.65 | | | | | | | |
| *Escherichia coli* str. K12 substr. MG1655 | 45.37 | 44.3 | 42.6 | 44.04 | 44.26 | 44.26 | 44.15 | 43.94 | 43.98 | 43.95 | 44.03 | 44.06 | 44.14 | 44.03 | 44.11 | | | | | | |
| *Gloeobacter violaceus* PCC 7421 | 42.39 | 55.01 | 54.66 | 55.22 | 56.05 | 56.22 | 55.88 | 56.07 | 55.39 | 55.43 | 55.44 | 55.37 | 55.4 | 55.52 | 55.23 | 44.19 | | | | | |
| *Nostoc azollae* 0708 | 42.06 | 78.24 | 77.17 | 78.14 | 75.68 | 75.86 | 75.91 | 78.24 | 77.76 | 77.76 | 77.73 | 77.83 | 77.81 | 77.76 | 78.27 | 44.27 | 55.93 | | | | |
| *Nostoc carneum* NIES 2107 | 43.46 | 73.02 | 73.8 | 73.48 | 76.55 | 76.36 | 95.21 | 76.62 | 73.91 | 73.89 | 73.9 | 73.88 | 73.92 | 73.72 | 73.85 | 44.23 | 55.7 | 75.53 | | | |
| *Nostoc sphaeroides* strain Kutzing | 41.78 | 73.87 | 73.04 | 74.02 | 77.57 | 77.48 | 77.82 | 78.94 | 74.2 | 74.18 | 74.21 | 74.3 | 74.16 | 74.3 | 74.39 | 44.13 | 56.07 | 76.5 | 77.4 | | |
| *Scytonema crispum* CAWBG524 | 42.86 | 72.72 | 73.04 | 72.63 | 75.6 | 75.6 | 76.15 | 76.43 | 73.45 | 73.43 | 73.49 | 73.58 | 73.3 | 73.35 | 73.24 | 44.08 | 56.54 | 74.82 | 75.79 | 76.98 | |
| *Scytonema crispum* CAWBG72 | 43.18 | 72.91 | 73.39 | 72.91 | 75.57 | 75.5 | 75.86 | 76.31 | 73.7 | 73.68 | 73.76 | 73.83 | 73.49 | 73.5 | 73.46 | 44.02 | 56.52 | 74.88 | 75.7 | 76.95 | 99.12 |

**Taxonomic Cutoffs**

- 95% - 100%; Species
- 65% - 95%; Genus
- 45% - 65%; Family
- ≤45%; Order

**Figure S2**: Triangle heatmap of pairwise AAI calculations. All numbers in the heatmap are listed as percentages. The four genomes from this study are labelled in bold. Colours indicate the taxonomic identity standards based on AAI as described by Konstantinidis et al. (2017).

#### 3 Abundance of protein coding genes

The abundance profile of protein coding genes within each strain (ACBU01, ACMB03, ACMB13 and FSS-124) was performed using protein and molecular function annotation databases to classify genes. The databases used were: Cluster of Orthlogous Groups of proteins (COGs) (Tatusov et al., 2000), KEGG Orthology (KO) (Kanehisa and Goto, 2000), Pfam and TIGRfam. Estimated gene copies were used to produce matrices with counts for all functions in a database for all strains. Matrices were filtered for functions found only in one or more strains but not all strains.

**Supplementary Table S2**: Strain-specific protein coding genes based on COGs.

| Function ID | Function name | ACBU01 | ACMB03 | ACMB13 | FSS-124 |
|---|---|---|---|---|---|
| **ACBU01 only** | | | | | |
| COG1813 | Archaeal ribosome-binding protein aMBF1, putative translation factor, contains Zn-ribbon and HTH domains | 1 | 0 | 0 | 0 |
| COG3312 | FoF1-type ATP synthase assembly protein I | 2 | 0 | 0 | 0 |
| **ACMB13 only** | | | | | |
| COG1326 | Uncharacterized archaeal Zn-finger protein | 0 | 0 | 1 | 0 |
| **FSS-124 only** | | | | | |
| COG0011 | Uncharacterized conserved protein YqgV, UPF0045/DUF77 family | 0 | 0 | 0 | 2 |
| COG0267 | Ribosomal protein L33 | 0 | 0 | 0 | 1 |
| COG0659 | Sulfate permease or related transporter, MFS superfamily | 0 | 0 | 0 | 1 |
| COG0732 | Restriction endonuclease S subunit | 0 | 0 | 0 | 3 |
| COG0753 | Catalase | 0 | 0 | 0 | 1 |
| COG1030 | Membrane-bound serine protease (ClpP class) | 0 | 0 | 0 | 1 |
| COG1240 | Mg-chelatase subunit ChlD | 0 | 0 | 0 | 2 |
| COG1269 | Archaeal/vacuolar-type H+-ATPase subunit I/STV1 | 0 | 0 | 0 | 1 |
| COG1340 | Uncharacterized coiled-coil protein, contains DUF342 domain | 0 | 0 | 0 | 1 |
| COG1360 | Flagellar motor protein MotB | 0 | 0 | 0 | 2 |
| COG1382 | Prefoldin, chaperonin cofactor | 0 | 0 | 0 | 2 |
| COG1491 | Predicted nucleic acid-binding OB-fold protein | 0 | 0 | 0 | 1 |
| COG1566 | Multidrug resistance efflux pump | 0 | 0 | 0 | 2 |
| COG1633 | Rubrerythrin | 0 | 0 | 0 | 3 |
| COG1648 | Siroheme synthase (precorrin-2 oxidase/ferrochelatase domain) | 0 | 0 | 0 | 1 |
| COG1694 | NTP pyrophosphatase, house-cleaning of non-canonical NTPs | 0 | 0 | 0 | 1 |
| COG1811 | Uncharacterized membrane protein YqgA, affects biofilm formation | 0 | 0 | 0 | 1 |
| COG1843 | Flagellar hook assembly protein FlgD | 0 | 0 | 0 | 1 |
| COG1859 | RNA:NAD 2'-phosphotransferase, TPT1/KptA family | 0 | 0 | 0 | 1 |
| COG1861 | Spore coat polysaccharide biosynthesis protein SpsF, cytidylyltransferase family | 0 | 0 | 0 | 1 |

| Function ID | Function name | ACBU01 | ACMB03 | ACMB13 | FSS-124 |
|---|---|---|---|---|---|
| COG1895 | Uncharacterised protein, contains HEPN domain, UPF0332 family | 0 | 0 | 0 | 3 |
| COG2019 | Archaeal adenylate kinase | 0 | 0 | 0 | 1 |
| COG2020 | Protein-S-isoprenylcysteine O-methyltransferase Ste14 | 0 | 0 | 0 | 1 |
| COG2028 | Uncharacterized protein | 0 | 0 | 0 | 1 |
| COG2089 | Sialic acid synthase SpsE, contains C-terminal SAF domain | 0 | 0 | 0 | 1 |
| COG2256 | Replication-associated recombination protein RarA (DNA-dependent ATPase) | 0 | 0 | 0 | 1 |
| COG2303 | Choline dehydrogenase or related flavoprotein | 0 | 0 | 0 | 1 |
| COG2379 | Glycerate-2-kinase | 0 | 0 | 0 | 1 |
| COG2384 | tRNA A22 N-methylase | 0 | 0 | 0 | 1 |
| COG2426 | Uncharacterized membrane protein | 0 | 0 | 0 | 1 |
| COG2522 | Predicted transcriptional regulator | 0 | 0 | 0 | 1 |
| COG2770 | HAMP domain | 0 | 0 | 0 | 1 |
| COG2801 | Transposase InsO and inactivated derivatives | 0 | 0 | 0 | 1 |
| COG2900 | Uncharacterized coiled-coil protein SlyX (sensitive to lysis X) | 0 | 0 | 0 | 1 |
| COG2976 | Putative negative regulator of RcsB-dependent stress response | 0 | 0 | 0 | 2 |
| COG3025 | Inorganic triphosphatase YgiF, contains CYTH and CHAD domains | 0 | 0 | 0 | 1 |
| COG3064 | Membrane protein involved in colicin uptake | 0 | 0 | 0 | 1 |
| COG3113 | ABC-type transporter Mla maintaining outer membrane lipid asymmetry, MlaB component, contains STAS domain | 0 | 0 | 0 | 1 |
| COG3121 | P pilus assembly protein, chaperone PapD | 0 | 0 | 0 | 1 |
| COG3162 | Uncharacterized membrane protein, DUF485 family | 0 | 0 | 0 | 1 |
| COG3188 | Outer membrane usher protein FimD/PapC | 0 | 0 | 0 | 1 |
| COG3209 | Uncharacterized conserved protein RhaS, contains 28 RHS repeats | 0 | 0 | 0 | 9 |
| COG3275 | Sensor histidine kinase, LytS/YehU family | 0 | 0 | 0 | 1 |
| COG3311 | Predicted DNA-binding transcriptional regulator AlpA | 0 | 0 | 0 | 1 |
| COG3370 | Uncharacterized protein | 0 | 0 | 0 | 1 |
| COG3378 | Phage- or plasmid-associated DNA primase | 0 | 0 | 0 | 1 |
| COG3391 | DNA-binding beta-propeller fold protein YncE | 0 | 0 | 0 | 2 |

| Function ID | Function name | ACBU01 | ACMB03 | ACMB13 | FSS-124 |
|---|---|---|---|---|---|
| COG3432 | Predicted transcriptional regulator | 0 | 0 | 0 | 3 |
| COG3508 | Homogentisate 1,2-dioxygenase | 0 | 0 | 0 | 1 |
| COG3607 | Predicted lactoylglutathione lyase | 0 | 0 | 0 | 1 |
| COG3620 | Predicted transcriptional regulator with C-terminal CBS domains | 0 | 0 | 0 | 2 |
| COG3624 | Alpha-D-ribose 1-methylphosphonate 5-triphosphate synthase subunit PhnG | 0 | 0 | 0 | 1 |
| COG3636 | DNA-binding prophage protein | 0 | 0 | 0 | 1 |
| COG3651 | Uncharacterized conserved protein, DUF2237 family | 0 | 0 | 0 | 1 |
| COG3655 | DNA-binding transcriptional regulator, XRE family | 0 | 0 | 0 | 2 |
| COG3657 | Putative component of the toxin-antitoxin plasmid stabilization module | 0 | 0 | 0 | 1 |
| COG3673 | Uncharacterized protein, PA2063/DUF2235 family | 0 | 0 | 0 | 2 |
| COG3742 | Uncharacterized protein, contains PIN domain | 0 | 0 | 0 | 1 |
| COG3751 | Proline 4-hydroxylase (includes Rps23 Pro-64 3,4-dihydroxylase Tpa1), contains SM-20 domain | 0 | 0 | 0 | 1 |
| COG3752 | Steroid 5-alpha reductase family enzyme | 0 | 0 | 0 | 1 |
| COG3883 | Uncharacterized N-terminal domain of peptidoglycan hydrolase CwlO | 0 | 0 | 0 | 1 |
| COG3906 | Uncharacterized protein YrzB, UPF0473 family | 0 | 0 | 0 | 1 |
| COG3910 | Predicted ATPase | 0 | 0 | 0 | 2 |
| COG3980 | Spore coat polysaccharide biosynthesis protein SpsG, predicted glycosyltransferase | 0 | 0 | 0 | 1 |
| COG4020 | Uncharacterized protein | 0 | 0 | 0 | 1 |
| COG4022 | Uncharacterized protein | 0 | 0 | 0 | 1 |
| COG4075 | Uncharacterized protein, distantly related to nitrogen regulatory protein PII | 0 | 0 | 0 | 1 |
| COG4293 | Uncharacterized protein | 0 | 0 | 0 | 1 |
| COG4304 | Uncharacterized protein | 0 | 0 | 0 | 1 |
| COG4333 | Uncharacterized protein | 0 | 0 | 0 | 1 |
| COG4568 | Transcriptional antiterminator Rof (Rho-off) | 0 | 0 | 0 | 1 |
| COG4589 | Predicted CDP-diglyceride synthetase/phosphatidate cytidylyltransferase | 0 | 0 | 0 | 1 |
| COG4645 | Uncharacterized protein | 0 | 0 | 0 | 1 |
| COG4681 | Uncharacterized conserved protein YaeQ, suppresses RfaH defect | 0 | 0 | 0 | 1 |

| Function ID | Function name | ACBU01 | ACMB03 | ACMB13 | FSS-124 |
|---|---|---|---|---|---|
| COG4683 | Uncharacterized protein | 0 | 0 | 0 | 3 |
| COG4687 | Uncharacterized protein | 0 | 0 | 0 | 2 |
| COG4874 | Uncharacterized protein | 0 | 0 | 0 | 1 |
| COG5002 | Signal transduction histidine kinase | 0 | 0 | 0 | 1 |
| COG5010 | Flp pilus assembly protein TadD, contains TPR repeats | 0 | 0 | 0 | 2 |
| COG5074 | t-SNARE complex subunit, syntaxin | 0 | 0 | 0 | 1 |
| COG5207 | Uncharacterized Zn-finger protein, UBP-type | 0 | 0 | 0 | 1 |
| COG5280 | Phage-related minor tail protein | 0 | 0 | 0 | 1 |
| COG5331 | Uncharacterized protein | 0 | 0 | 0 | 1 |
| COG5428 | Uncharacterized protein YuzE | 0 | 0 | 0 | 2 |
| COG5430 | Spore coat protein U (SCPU) domain, function unknown | 0 | 0 | 0 | 2 |
| COG5509 | Uncharacterized small protein, DUF1192 family | 0 | 0 | 0 | 2 |
| **ACBU01 and FSS-124 only** | | | | | |
| COG3599 | Cell division septum initiation DivIVA, interacts with FtsZ, MinD and other proteins | 2 | 0 | 0 | 2 |
| **ACMB03 and FSS-124 only** | | | | | |
| COG1236 | RNA processing exonuclease, beta-lactamase fold, Cft2 family | 0 | 1 | 0 | 1 |
| **ACBU01, ACMB03 and ACMB13 only** | | | | | |
| COG0218 | GTP-binding protein EngB required for normal cell division | 3 | 3 | 3 | 0 |
| COG0241 | Histidinol phosphatase or a related phosphatase | 1 | 1 | 1 | 0 |
| COG0271 | Stress-induced morphogen (activity unknown) | 1 | 1 | 1 | 0 |
| COG0279 | Phosphoheptose isomerase | 1 | 1 | 1 | 0 |
| COG0432 | Thiamin phosphate synthase YjbQ, UPF0047 family | 1 | 1 | 1 | 0 |
| COG0439 | Biotin carboxylase | 1 | 1 | 1 | 0 |
| COG0455 | MinD-like ATPase involved in chromosome partitioning or flagellar assembly | 1 | 1 | 1 | 0 |
| COG0492 | Thioredoxin reductase | 1 | 1 | 1 | 0 |
| COG0693 | Putative intracellular protease/amidase | 2 | 2 | 2 | 0 |

| Function ID | Function name | ACBU01 | ACMB03 | ACMB13 | FSS-124 |
|---|---|---|---|---|---|
| COG0710 | 3-dehydroquinate dehydratase | 1 | 1 | 1 | 0 |
| COG0737 | 2',3'-cyclic-nucleotide 2'-phosphodiesterase/5'- or 3'-nucleotidase, 5'-nucleotidase family | 1 | 1 | 1 | 0 |
| COG0786 | Na+/glutamate symporter | 1 | 1 | 1 | 0 |
| COG1317 | Flagellar biosynthesis/type III secretory pathway protein FliH | 1 | 1 | 1 | 0 |
| COG1427 | Predicted periplasmic solute-binding protein | 1 | 1 | 1 | 0 |
| COG1474 | Cdc6-related protein, AAA superfamily ATPase | 2 | 2 | 2 | 0 |
| COG1483 | Predicted ATPase, AAA+ superfamily | 1 | 1 | 1 | 0 |
| COG1652 | Nucleoid-associated protein YgaU, contains BON and LysM domains | 1 | 1 | 1 | 0 |
| COG1658 | 5S rRNA maturation endonuclease (Ribonuclease M5), contains TOPRIM domain | 1 | 1 | 1 | 0 |
| COG1730 | Prefoldin subunit 5 | 1 | 1 | 1 | 0 |
| COG1872 | Uncharacterized conserved protein YggU, UPF0235/DUF167 family | 1 | 1 | 1 | 0 |
| COG1917 | Cupin domain protein related to quercetin dioxygenase | 2 | 2 | 2 | 0 |
| COG1923 | sRNA-binding regulator protein Hfq | 1 | 1 | 1 | 0 |
| COG2189 | Adenine specific DNA methylase Mod | 2 | 2 | 2 | 0 |
| COG2413 | Predicted nucleotidyltransferase | 4 | 4 | 4 | 0 |
| COG2870 | ADP-heptose synthase, bifunctional sugar kinase/adenylyltransferase | 2 | 2 | 2 | 0 |
| COG2874 | Archaellum biogenesis protein FlaH, an ATPase | 1 | 1 | 1 | 0 |
| COG2880 | Predicted DNA-binding protein, potential antitoxin AbrB/MazE fold | 1 | 1 | 1 | 0 |
| COG2916 | DNA-binding protein H-NS | 1 | 1 | 1 | 0 |
| COG2923 | Sulfur relay (sulfurtransferase) complex TusC component, DsrF/TusC family | 1 | 1 | 1 | 0 |
| COG2942 | Mannose or cellobiose epimerase, N-acyl-D-glucosamine 2-epimerase family | 2 | 2 | 2 | 0 |
| COG3264 | Small-conductance mechanosensitive channel | 1 | 1 | 1 | 0 |
| COG3310 | Uncharacterized protein | 1 | 1 | 1 | 0 |
| COG3385 | IS4 transposase | 2 | 2 | 1 | 0 |
| COG3433 | Aryl carrier domain | 1 | 1 | 1 | 0 |
| COG3461 | Uncharacterized protein | 1 | 1 | 1 | 0 |
| COG3514 | Uncharacterized conserved protein, DUF4415 family | 1 | 1 | 1 | 0 |

| Function ID | Function name | ACBU01 | ACMB03 | ACMB13 | FSS-124 |
|---|---|---|---|---|---|
| COG3824 | Predicted Zn-dependent protease, minimal metalloprotease (MMP)-like domain | 1 | 1 | 1 | 0 |
| COG3861 | Stress response protein YsnF (function unknown) | 1 | 1 | 1 | 0 |
| COG3863 | Uncharacterized protein YycO | 1 | 1 | 1 | 0 |
| COG3905 | Predicted transcriptional regulator | 3 | 3 | 1 | 0 |
| COG3972 | Superfamily I DNA and RNA helicases | 2 | 2 | 2 | 0 |
| COG4785 | Lipoprotein NlpI, contains TPR repeats | 2 | 2 | 2 | 0 |
| COG4804 | Predicted nuclease of restriction endonuclease-like (RecB) superfamily, DUF1016 family | 1 | 1 | 1 | 0 |
| COG4850 | Phosphatidate phosphatase APP1 | 1 | 1 | 1 | 0 |
| COG4942 | Septal ring factor EnvC, activator of murein hydrolases AmiA and AmiB | 3 | 1 | 1 | 0 |
| COG5341 | Uncharacterized protein | 2 | 2 | 2 | 0 |
| COG5611 | Predicted nucleic-acid-binding protein, contains PIN domain | 1 | 1 | 1 | 0 |
| COG5619 | Uncharacterized protein | 1 | 1 | 1 | 0 |
| **ACMB03, ACMB13 and FSS-124 only** | | | | | |
| COG1511 | Uncharacterized membrane protein YhgE, phage infection protein (PIP) family | 0 | 2 | 2 | 1 |

**Supplementary Table S3**: Strain-specific protein coding genes based on KO. EC numbers refer to the Enzyme Nomenclature by KEGG.

| Function ID | Function name | ACBU01 | ACMB03 | ACMB13 | FSS-124 |
|---|---|---|---|---|---|
| **FSS-124 only** | | | | | |
| KO:K00694 | cellulose synthase (UDP-forming) [EC:2.4.1.12] | 0 | 0 | 0 | 1 |
| KO:K01654 | N-acetylneuraminate synthase [EC:2.5.1.56] | 0 | 0 | 0 | 1 |
| KO:K02913 | large subunit ribosomal protein L33 | 0 | 0 | 0 | 1 |
| KO:K03335 | inosose dehydratase [EC:4.2.1.44] | 0 | 0 | 0 | 1 |
| KO:K06919 | putative DNA primase/helicase | 0 | 0 | 0 | 1 |
| KO:K06926 | uncharacterized protein | 0 | 0 | 0 | 1 |
| KO:K07000 | uncharacterized protein | 0 | 0 | 0 | 1 |
| KO:K07063 | uncharacterized protein | 0 | 0 | 0 | 1 |
| KO:K07346 | fimbrial chaperone protein | 0 | 0 | 0 | 1 |
| KO:K07347 | outer membrane usher protein | 0 | 0 | 0 | 1 |
| KO:K07454 | putative restriction endonuclease | 0 | 0 | 0 | 1 |
| KO:K07495 | putative transposase | 0 | 0 | 0 | 1 |
| KO:K07559 | putative RNA 2'-phosphotransferase [EC:2.7.1.-] | 0 | 0 | 0 | 1 |
| KO:K09966 | uncharacterized protein | 0 | 0 | 0 | 1 |
| KO:K11987 | prostaglandin-endoperoxide synthase 2 [EC:1.14.99.1] | 0 | 0 | 0 | 1 |
| KO:K16050 | 4,5:9,10-diseco-3-hydroxy-5,9,17-trioxoandrosta-1(10),2-diene-4-oate hydrolase [EC:3.7.1.17] | 0 | 0 | 0 | 1 |
| KO:K17062 | arginine/lysine/histidine/glutamine transport system substrate-binding and permease protein | 0 | 0 | 0 | 1 |
| KO:K17247 | methionine sulfoxide reductase heme-binding subunit | 0 | 0 | 0 | 1 |
| KO:K19465 | mitochondrial genome maintenance exonuclease 1 [EC:3.1.-.-] | 0 | 0 | 0 | 1 |
| KO:K19686 | ribonuclease VapC [EC:3.1.-.-] | 0 | 0 | 0 | 1 |
| KO:K19687 | antitoxin VapB | 0 | 0 | 0 | 1 |
| **ACBU01, ACMB03 and ACMB13 only** | | | | | |
| KO:K00613 | glycine amidinotransferase [EC:2.1.4.1] | 1 | 1 | 1 | 0 |
| KO:K00754 | - | 1 | 1 | 1 | 0 |
| KO:K01081 | 5'-nucleotidase [EC:3.1.3.5] | 1 | 1 | 1 | 0 |

| Function ID | Function name | ACBU01 | ACMB03 | ACMB13 | FSS-124 |
|---|---|---|---|---|---|
| KO:K01176 | alpha-amylase [EC:3.2.1.1] | 1 | 1 | 1 | 0 |
| KO:K01234 | neopullulanase [EC:3.2.1.135] | 1 | 1 | 1 | 0 |
| KO:K02805 | dTDP-4-amino-4,6-dideoxygalactose transaminase [EC:2.6.1.59] | 1 | 1 | 1 | 0 |
| KO:K03271 | D-sedoheptulose 7-phosphate isomerase [EC:5.3.1.28] | 1 | 1 | 1 | 0 |
| KO:K03273 | D-glycero-D-manno-heptose 1,7-bisphosphate phosphatase [EC:3.1.3.82 3.1.3.83] | 1 | 1 | 1 | 0 |
| KO:K03325 | arsenite transporter, ACR3 family | 1 | 1 | 1 | 0 |
| KO:K03427 | type I restriction enzyme M protein [EC:2.1.1.72] | 2 | 2 | 2 | 0 |
| KO:K07001 | NTE family protein | 1 | 1 | 1 | 0 |
| KO:K09131 | uncharacterized protein | 1 | 1 | 1 | 0 |
| KO:K16710 | colanic acid/amylovoran biosynthesis protein | 1 | 1 | 1 | 0 |
| KO:K18654 | kanosamine-6-phosphate phosphatase [EC:3.1.3.92] | 1 | 1 | 1 | 0 |
| KO:K19169 | DNA sulfur modification protein DndB | 1 | 1 | 1 | 0 |
| KO:K19429 | acetyltransferase EpsM [EC:2.3.1.-] | 1 | 1 | 1 | 0 |
| KO:K19547 | bacilysin biosynthesis protein BacB [EC:5.-.-.-] | 1 | 1 | 1 | 0 |
| KO:K19550 | bacilysin biosynthesis oxidoreductase BacG [EC:1.-.-.-] | 1 | 1 | 1 | 0 |
| **ACBU01, ACMB03 and FSS-124 only** | | | | | |
| KO:K02662 | type IV pilus assembly protein PilM | 1 | 1 | 0 | 1 |
| **ACBU01, ACMB13 and FSS-124 only** | | | | | |
| KO:K02650 | type IV pilus assembly protein PilA | 1 | 0 | 1 | 1 |
| **ACMB03, ACMB13 and FSS-124 only** | | | | | |
| KO:K02666 | type IV pilus assembly protein PilQ | 0 | 1 | 1 | 1 |

**Supplementary Table S4**: Strain-specific protein coding genes based on Pfam.

| Function ID | Function name | ACBU01 | ACMB03 | ACMB13 | FSS-124 |
|---|---|---|---|---|---|
| **FSS-124 only** | | | | | |
| pfam00028 | Cadherin domain | 0 | 0 | 0 | 1 |
| pfam00045 | Hemopexin | 0 | 0 | 0 | 3 |
| pfam00139 | Legume lectin domain | 0 | 0 | 0 | 1 |
| pfam00151 | Lipase | 0 | 0 | 0 | 1 |
| pfam00345 | Pili and flagellar-assembly chaperone, PapD N-terminal domain | 0 | 0 | 0 | 1 |
| pfam00471 | Ribosomal protein L33 | 0 | 0 | 0 | 1 |
| pfam00577 | Outer membrane usher protein | 0 | 0 | 0 | 1 |
| pfam00665 | Integrase core domain | 0 | 0 | 0 | 1 |
| pfam01420 | Type I restriction modification DNA specificity domain | 0 | 0 | 0 | 2 |
| pfam01569 | PAP2 superfamily | 0 | 0 | 0 | 1 |
| pfam01764 | Lipase (class 3) | 0 | 0 | 0 | 1 |
| pfam01885 | RNA 2'-phosphotransferase, Tpt1 / KptA family | 0 | 0 | 0 | 1 |
| pfam02348 | Cytidylyltransferase | 0 | 0 | 0 | 1 |
| pfam03098 | Animal haem peroxidase | 0 | 0 | 0 | 1 |
| pfam03102 | NeuB family | 0 | 0 | 0 | 1 |
| pfam03552 | Cellulose synthase | 0 | 0 | 0 | 2 |
| pfam03747 | ADP-ribosylglycohydrolase | 0 | 0 | 0 | 1 |
| pfam04140 | Isoprenylcysteine carboxyl methyltransferase (ICMT) family | 0 | 0 | 0 | 1 |
| pfam04261 | Dyp-type peroxidase family | 0 | 0 | 0 | 1 |
| pfam04505 | Interferon-induced transmembrane protein | 0 | 0 | 0 | 1 |
| pfam04989 | Cephalosporin hydroxylase | 0 | 0 | 0 | 1 |
| pfam05199 | GMC oxidoreductase | 0 | 0 | 0 | 1 |
| pfam05229 | Spore Coat Protein U domain | 0 | 0 | 0 | 2 |
| pfam05593 | RHS Repeat | 0 | 0 | 0 | 10 |
| pfam05728 | Uncharacterised protein family (UPF0227) | 0 | 0 | 0 | 1 |

| Function ID | Function name | ACBU01 | ACMB03 | ACMB13 | FSS-124 |
|---|---|---|---|---|---|
| pfam06695 | Putative small multi-drug export protein | 0 | 0 | 0 | 1 |
| pfam07799 | Protein of unknown function (DUF1643) | 0 | 0 | 0 | 1 |
| pfam08417 | Pheophorbide a oxygenase | 0 | 0 | 0 | 1 |
| pfam08666 | SAF domain | 0 | 0 | 0 | 1 |
| pfam08706 | D5 N terminal like | 0 | 0 | 0 | 1 |
| pfam08797 | HIRAN domain | 0 | 0 | 0 | 1 |
| pfam08937 | MTH538 TIR-like domain (DUF1863) | 0 | 0 | 0 | 1 |
| pfam08972 | Domain of unknown function (DUF1902) | 0 | 0 | 0 | 1 |
| pfam09564 | NgoBV restriction endonuclease | 0 | 0 | 0 | 1 |
| pfam09957 | Bacterial antitoxin of type II TA system, VapB | 0 | 0 | 0 | 1 |
| pfam09994 | Uncharacterized alpha/beta hydrolase domain (DUF2235) | 0 | 0 | 0 | 1 |
| pfam09996 | Uncharacterized protein conserved in bacteria (DUF2237) | 0 | 0 | 0 | 1 |
| pfam10049 | Protein of unknown function (DUF2283) | 0 | 0 | 0 | 2 |
| pfam10544 | T5orf172 domain | 0 | 0 | 0 | 1 |
| pfam10825 | Protein of unknown function (DUF2752) | 0 | 0 | 0 | 1 |
| pfam11583 | P-aminobenzoate N-oxygenase AurF | 0 | 0 | 0 | 1 |
| pfam11867 | Domain of unknown function (DUF3387) | 0 | 0 | 0 | 1 |
| pfam12686 | Protein of unknown function (DUF3800) | 0 | 0 | 0 | 1 |
| pfam12872 | OST-HTH/LOTUS domain | 0 | 0 | 0 | 1 |
| pfam12902 | Ferritin-like | 0 | 0 | 0 | 1 |
| pfam13205 | Bacterial Ig-like domain | 0 | 0 | 0 | 1 |
| pfam13207 | AAA domain | 0 | 0 | 0 | 1 |
| pfam13246 | Cation transport ATPase (P-type) | 0 | 0 | 0 | 1 |
| pfam13385 | Concanavalin A-like lectin/glucanases superfamily | 0 | 0 | 0 | 1 |
| pfam13420 | Acetyltransferase (GNAT) domain | 0 | 0 | 0 | 1 |
| pfam13448 | Domain of unknown function (DUF4114) | 0 | 0 | 0 | 2 |
| pfam13578 | Methyltransferase domain | 0 | 0 | 0 | 1 |

| Function ID | Function name | ACBU01 | ACMB03 | ACMB13 | FSS-124 |
|---|---|---|---|---|---|
| pfam13860 | FlgD Ig-like domain | 0 | 0 | 0 | 1 |
| pfam14022 | Protein of unknown function (DUF4238) | 0 | 0 | 0 | 1 |
| pfam14355 | Abortive infection C-terminus | 0 | 0 | 0 | 1 |
| pfam14424 | The BURPS668_1122 family of deaminases | 0 | 0 | 0 | 1 |
| pfam14568 | SMI1-KNR4 cell-wall | 0 | 0 | 0 | 1 |
| pfam14938 | Soluble NSF attachment protein, SNAP | 0 | 0 | 0 | 1 |
| pfam16198 | tRNA pseudouridylate synthase B C-terminal domain | 0 | 0 | 0 | 1 |
| pfam16203 | ERCC3/RAD25/XPB C-terminal helicase | 0 | 0 | 0 | 1 |
| **ACBU01, ACMB03 and ACMB13 only** | | | | | |
| pfam00313 | 'Cold-shock' DNA-binding domain | 1 | 1 | 1 | 0 |
| pfam00376 | MerR family regulatory protein | 1 | 1 | 1 | 0 |
| pfam00668 | Condensation domain | 4 | 4 | 4 | 0 |
| pfam01535 | PPR repeat | 1 | 1 | 1 | 0 |
| pfam01609 | Transposase DDE domain | 2 | 2 | 1 | 0 |
| pfam01894 | Uncharacterised protein family UPF0047 | 1 | 1 | 1 | 0 |
| pfam01954 | Protein of unknown function DUF104 | 1 | 1 | 1 | 0 |
| pfam01965 | DJ-1/PfpI family | 1 | 1 | 1 | 0 |
| pfam02594 | Uncharacterised ACR, YggU family COG1872 | 1 | 1 | 1 | 0 |
| pfam02872 | 5'-nucleotidase, C-terminal domain | 1 | 1 | 1 | 0 |
| pfam02963 | Restriction endonuclease EcoRI | 1 | 1 | 1 | 0 |
| pfam03567 | Sulfotransferase family | 1 | 1 | 1 | 0 |
| pfam04266 | ASCH domain | 1 | 1 | 1 | 0 |
| pfam04465 | Protein of unknown function (DUF499) | 1 | 1 | 1 | 0 |
| pfam04577 | Protein of unknown function (DUF563) | 1 | 1 | 1 | 0 |
| pfam06250 | Protein of unknown function (DUF1016) | 1 | 1 | 1 | 0 |
| pfam08662 | Eukaryotic translation initiation factor eIF2A | 1 | 1 | 1 | 0 |
| pfam09519 | HindVP restriction endonuclease | 1 | 1 | 1 | 0 |

| Function ID | Function name | ACBU01 | ACMB03 | ACMB13 | FSS-124 |
|---|---|---|---|---|---|
| pfam09520 | Type II restriction endonuclease, TdeIII | 1 | 1 | 1 | 0 |
| pfam09557 | Domain of unknown function (DUF2382) | 1 | 1 | 1 | 0 |
| pfam10053 | Uncharacterized conserved protein (DUF2290) | 1 | 1 | 1 | 0 |
| pfam11845 | Protein of unknown function (DUF3365) | 1 | 1 | 1 | 0 |
| pfam12740 | Chlorophyllase enzyme | 1 | 1 | 1 | 0 |
| pfam12787 | EcsC protein family | 1 | 1 | 1 | 0 |
| pfam12973 | ChrR Cupin-like domain | 1 | 1 | 1 | 0 |
| pfam13000 | Acetyl-coenzyme A transporter 1 | 1 | 1 | 1 | 0 |
| pfam13245 | AAA domain | 1 | 1 | 1 | 0 |
| pfam13440 | Polysaccharide biosynthesis protein | 1 | 1 | 1 | 0 |
| pfam13516 | Leucine Rich repeat | 2 | 2 | 2 | 0 |
| pfam13535 | ATP-grasp domain | 1 | 1 | 1 | 0 |
| pfam13538 | UvrD-like helicase C-terminal domain | 2 | 2 | 2 | 0 |
| pfam13613 | Helix-turn-helix of DDE superfamily endonuclease | 1 | 1 | 1 | 0 |
| pfam13650 | Aspartyl protease | 3 | 3 | 3 | 0 |
| pfam13651 | Adenine-specific methyltransferase EcoRI | 1 | 1 | 1 | 0 |
| pfam14082 | Domain of unknown function (DUF4263) | 1 | 1 | 1 | 0 |
| pfam14384 | BrnA antitoxin of type II toxin-antitoxin system | 1 | 1 | 1 | 0 |
| pfam14524 | Wzt C-terminal domain | 1 | 1 | 1 | 0 |
| pfam14897 | EpsG family | 1 | 1 | 1 | 0 |
| pfam14903 | WG containing repeat | 9 | 9 | 9 | 0 |
| pfam16881 | N-terminal domain of lipoyl synthase of Radical_SAM family | 1 | 1 | 1 | 0 |
| **ACBU01, ACMB13 and FSS-124 only** | | | | | |
| pfam16734 | Type IV pilin-like G and H, putative | 1 | 0 | 1 | 1 |
| **ACMB03, ACMB13 and FSS-124 only** | | | | | |
| pfam03958 | Bacterial type II/III secretion system short domain | 0 | 1 | 1 | 1 |

| Function ID | Function name | ACBU01 | ACMB03 | ACMB13 | FSS-124 |
| --- | --- | --- | --- | --- | --- |
| pfam07660 | Secretin and TonB N terminus short domain | 0 | 1 | 1 | 1 |

**Supplementary Table S5**: Strain-specific protein coding genes based on TIGRfam.

| Function ID | Function name | ACBU01 | ACMB03 | ACMB13 | FSS-124 |
|---|---|---|---|---|---|
| **FSS-124 only** | | | | | |
| TIGR00348 | type I site-specific deoxyribonuclease, HsdR family | 0 | 0 | 0 | 1 |
| TIGR00358 | VacB and RNase II family 3'-5' exoribonucleases | 0 | 0 | 0 | 1 |
| TIGR00901 | AmpG-like permease | 0 | 0 | 0 | 1 |
| TIGR01023 | ribosomal protein L33, bacterial type | 0 | 0 | 0 | 1 |
| TIGR01230 | agmatinase | 0 | 0 | 0 | 1 |
| TIGR01523 | potassium and/or sodium efflux P-type ATPase, fungal-type | 0 | 0 | 0 | 1 |
| TIGR01643 | YD repeat (two copies) | 0 | 0 | 0 | 15 |
| TIGR01730 | RND family efflux transporter, MFP subunit | 0 | 0 | 0 | 1 |
| TIGR02081 | methionine biosynthesis protein MetW | 0 | 0 | 0 | 1 |
| TIGR02609 | putative addiction module antidote | 0 | 0 | 0 | 1 |
| TIGR02683 | putative addiction module killer protein | 0 | 0 | 0 | 1 |
| TIGR03174 | CRISPR type I-D/CYANO-associated protein Csc3/Cas10d | 0 | 0 | 0 | 1 |
| TIGR03585 | UDP-4-amino-4,6-dideoxy-N-acetyl-beta-L-altrosamine N-acetyltransferase | 0 | 0 | 0 | 1 |
| TIGR03586 | pseudaminic acid synthase | 0 | 0 | 0 | 1 |
| TIGR03588 | UDP-4-amino-4,6-dideoxy-N-acetyl-beta-L-altrosamine transaminase | 0 | 0 | 0 | 1 |
| TIGR03589 | UDP-N-acetylglucosamine 4,6-dehydratase (inverting) | 0 | 0 | 0 | 1 |
| TIGR03590 | UDP-2,4-diacetamido-2,4,6-trideoxy-beta-L-altropyranose hydrolase | 0 | 0 | 0 | 1 |
| TIGR03696 | RHS repeat-associated core domain | 0 | 0 | 0 | 1 |
| TIGR03763 | cyanoexosortase A | 0 | 0 | 0 | 1 |
| TIGR03798 | nif11-like leader peptide domain | 0 | 0 | 0 | 1 |
| TIGR04103 | nif11-class peptide radical SAM maturase 3 | 0 | 0 | 0 | 1 |
| TIGR04153 | cyanosortase A-associated protein | 0 | 0 | 0 | 1 |
| TIGR04379 | myo-inosose-2 dehydratase | 0 | 0 | 0 | 1 |
| TIGR04446 | prenylated cyclic peptide, anacyclamide/piricyclamide family | 0 | 0 | 0 | 1 |
| **ACBU01, ACMB03 and ACMB13 only** | | | | | |

| Function ID | Function name | ACBU01 | ACMB03 | ACMB13 | FSS-124 |
|---|---|---|---|---|---|
| TIGR00149 | secondary thiamine-phosphate synthase enzyme | 1 | 1 | 1 | 0 |
| TIGR00213 | D,D-heptose 1,7-bisphosphate phosphatase | 1 | 1 | 1 | 0 |
| TIGR00251 | TIGR00251 family protein | 1 | 1 | 1 | 0 |
| TIGR01409 | Tat (twin-arginine translocation) pathway signal sequence | 1 | 1 | 1 | 0 |
| TIGR01484 | HAD-superfamily hydrolase, subfamily IIB | 2 | 2 | 2 | 0 |
| TIGR02063 | ribonuclease R | 1 | 1 | 1 | 0 |
| TIGR02199 | rfaE bifunctional protein, domain II | 1 | 1 | 1 | 0 |
| TIGR02271 | conserved domain | 1 | 1 | 1 | 0 |
| TIGR04220 | cyanobactin biosynthesis protein, PatB/AcyB/McaB family | 1 | 1 | 1 | 0 |
| TIGR04447 | cyanobactin cluster PatC/TenC/TruC protein | 1 | 1 | 1 | 0 |
| TIGR04474 | three-Cys-motif partner protein | 1 | 1 | 1 | 0 |
| **ACBU01, ACMB03 and FSS-124 only** | | | | | |
| TIGR03438 | dimethylhistidine N-methyltransferase | 1 | 1 | 0 | 1 |

### 4 Analysis of genomic synteny

The alignment of genomes to infer synteny was performed on the IMG/MER system as described in the Materials and Methods. Specifically, the program MUMmer (Kurtz et al., 2004) was used to generate dotplot diagrams, using the input DNA sequences directly for comparing genomes with similar sequences (NUCmer). Vertical (green) and horizontal (brown) lines correspond to the scaffolds within each genome.

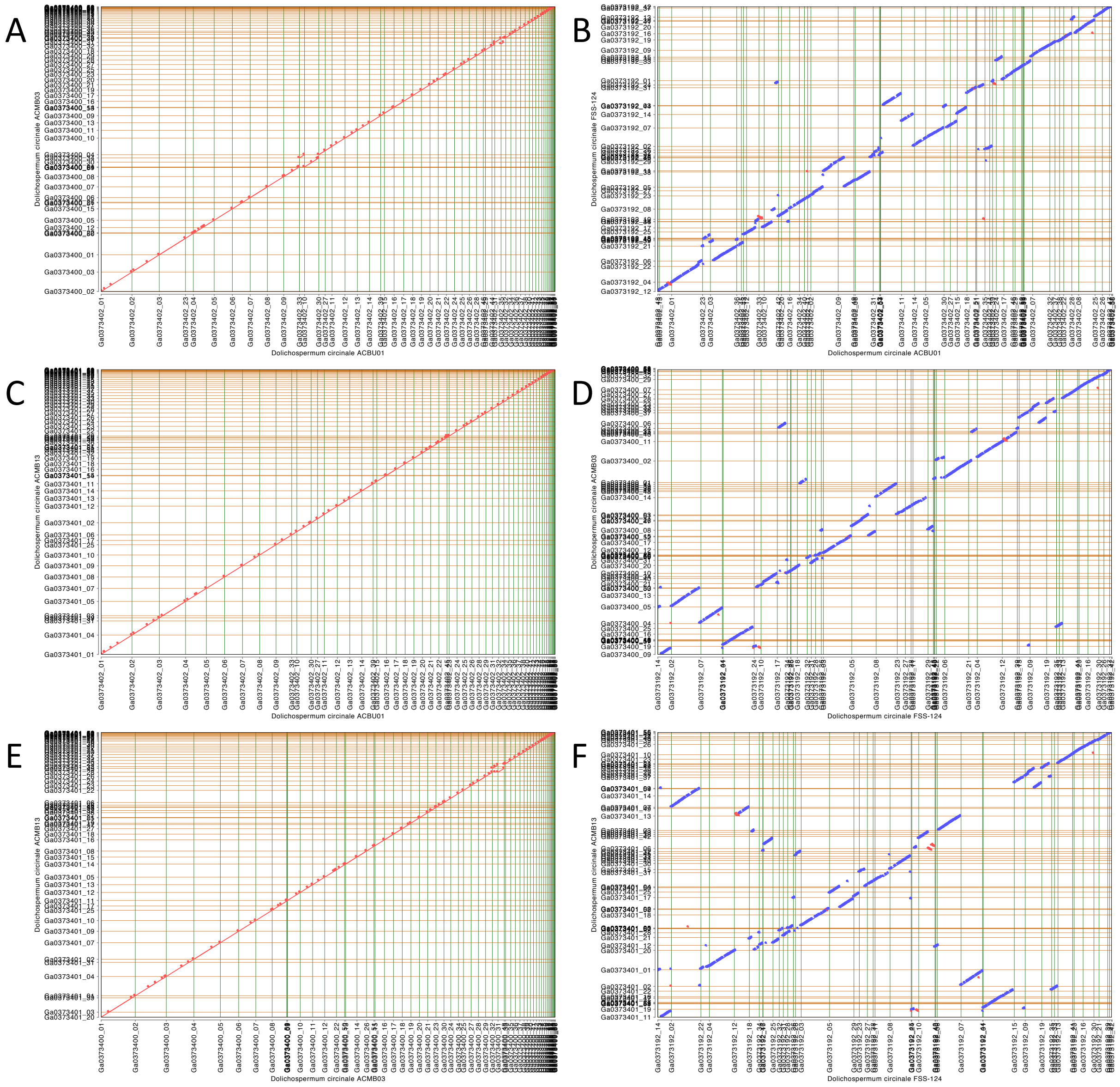


**Figure S3**: Synteny dotplots between the four *D. circinale* genomes: (A) ACBU01 and ACMB03, (B) ACBU01 and FSS-124, (C) ACBU01 and ACMB13, (D) FSS-124 and ACMB03, (E) ACMB03 and ACMB13, (F) FSS-124 and ACMB13.

#### 5 Identification of secondary metabolite clusters in *D. circinale*

The identification, annotation and analysis of secondary metabolite biosynthesis gene clusters present in all *D. circinale* genomes was performed using antiSMASH software version 5.0 (Blin et al., 2019) as described in the Materials and Methods. Three clusters with >80% similarity to known natural product clusters were found in all *D. circinale* genomes: geosmin synthase (Figure S4), anacyclamide (Figure S5) and a heterocyst glycolipid biosynthesis cluster (Figure S6). The figures show the genes in the query *D. circinale* genomes homologous to the annotated secondary metabolite biosynthetic genes. Annotations used the Minimum Information about a Biosynthetic Gene cluster (MIBiG) data standard (Medema et al., 2015). The MIBiG accession number (BCG#######) is provided. The chemical structures of identified compounds are present where relevant.

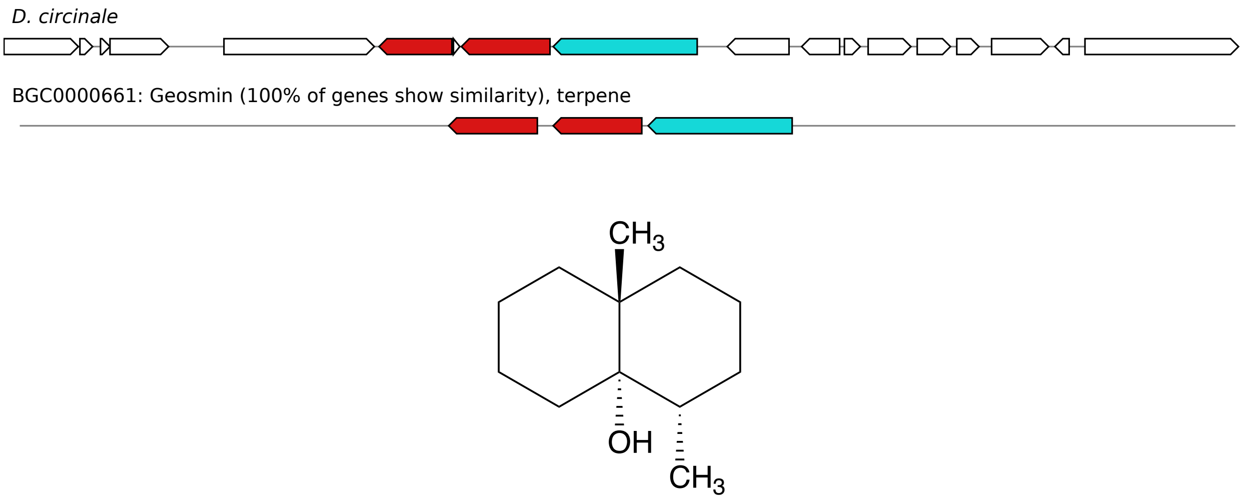


**Figure S4:** The geosmin synthase biosynthetic cluster identified in *D. circinale*, producing the organic compound geosmin.

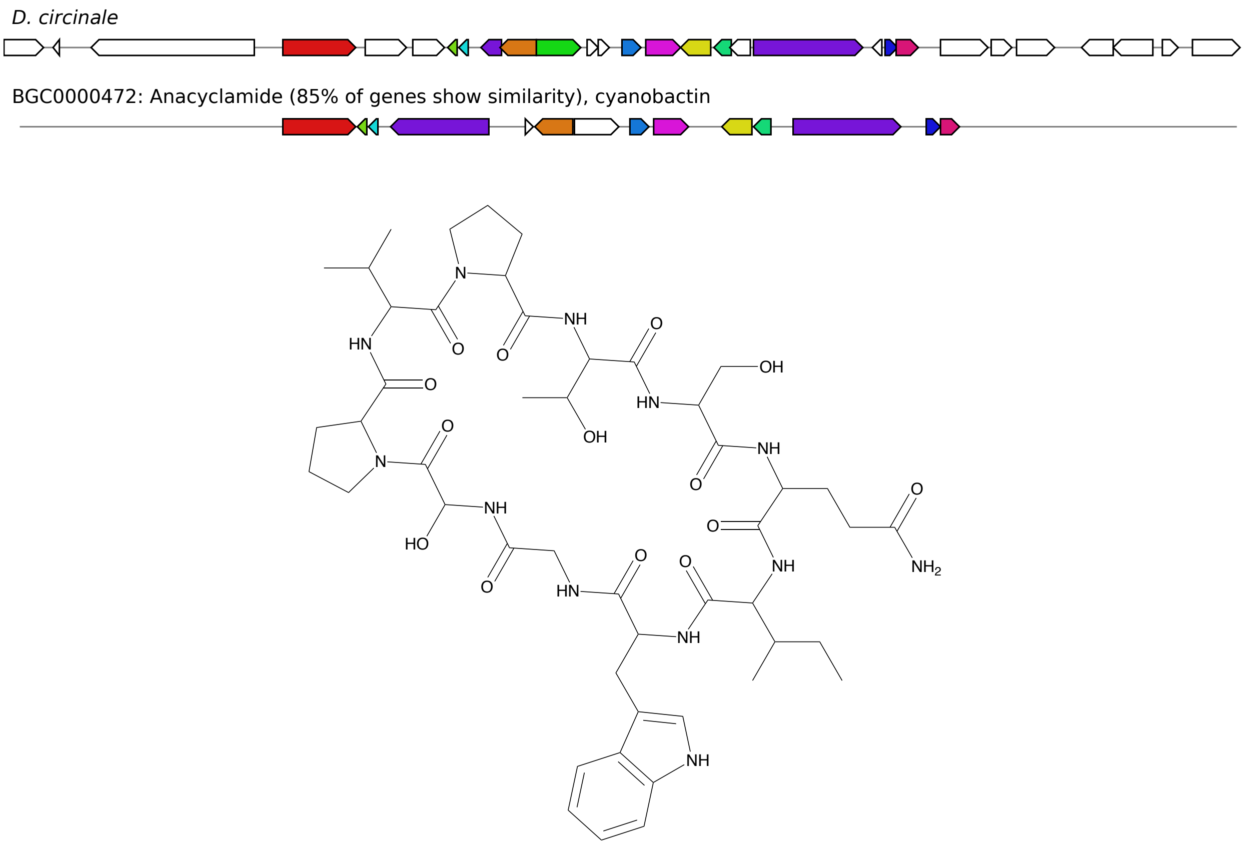


**Figure S5:** The anacyclamide biosynthesis cluster identified in *D. circinale*, producing the cyanobactin anacyclamide.

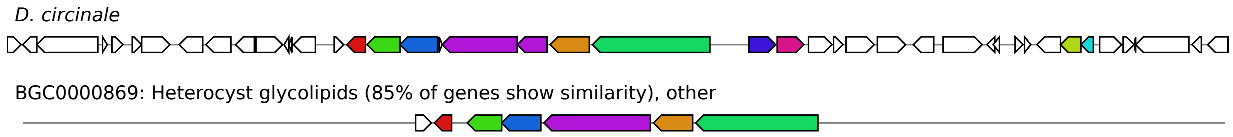


**Figure S6:** The heterocyst glycolipid biosynthesis cluster identified in *D. circinale*.

#### 6 Identification of *sxt* genes by PCR

Genomic DNA was extracted from the four *D. circinale* strains in this study and PCR was performed as detailed in the Materials and Methods. Gel electrophoresis of PCR products had expected amplicon sizes as follows: 3,191 bp using Acir_SUL_F and Acir_sxtI_R (Figure S7); 7,559 bp using Acir_SUL_F and Acir_SMF_R (Figure S8); 7,426 bp using Acir_sxtG_F and Acir_sxtI_R (Figure S9). L stands for the DNA ladder used in all gels (2-Log DNA ladder, New England BioLabs). The strains ACBU01, FSS-124, ACMB03 and ACMB13 are labelled 1-4 respectively.

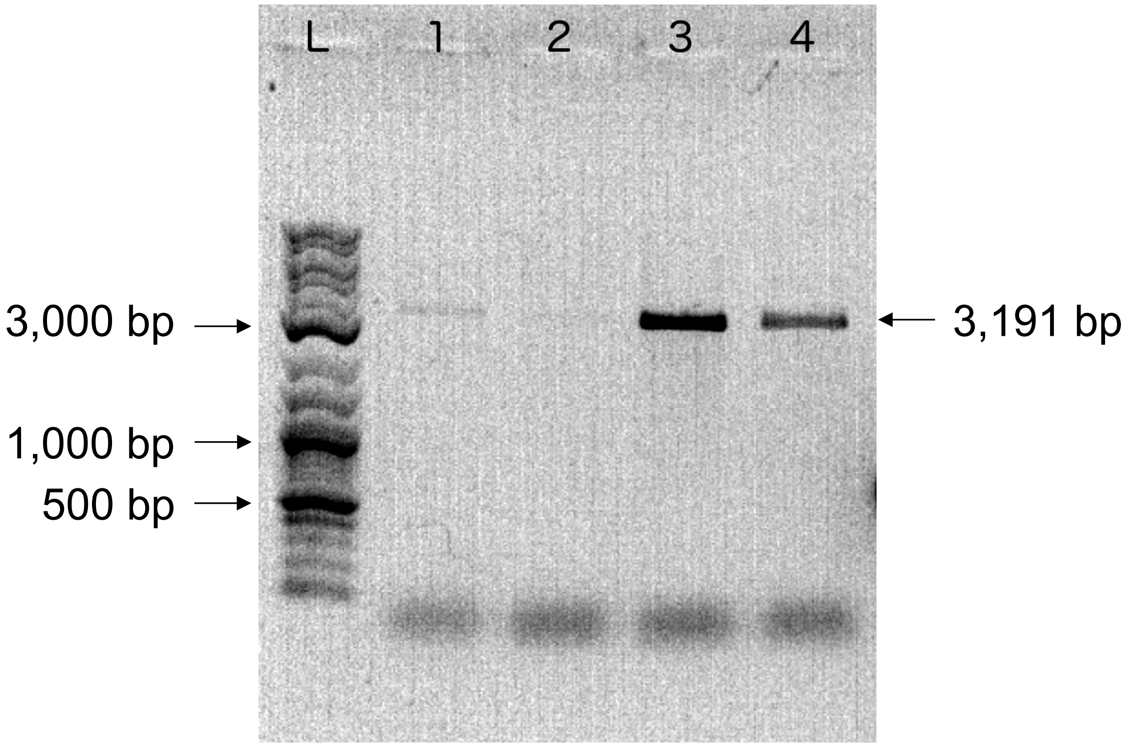


**Figure S7:** PCR amplicons generated from *D. circinale* genomic DNA using primers Acir_SUL_F and Acir_sxtI_R targeting the region between *sxtSUL* and *sxtI* in the *sxt* gene cluster.

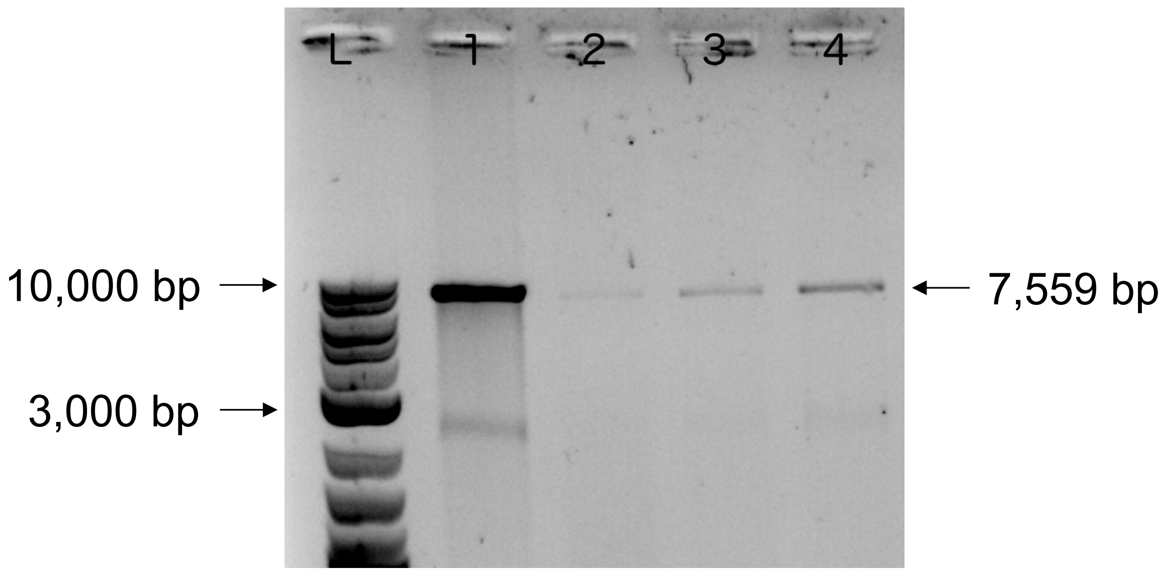


**Figure S8:** PCR amplicons generated from *D. circinale* genomic DNA using primers Acir_SUL_F and Acir_SMF_R, targeting the region between *sxtSUL* and the *smf* gene homolog at the 3'-end of the *sxt* gene cluster..

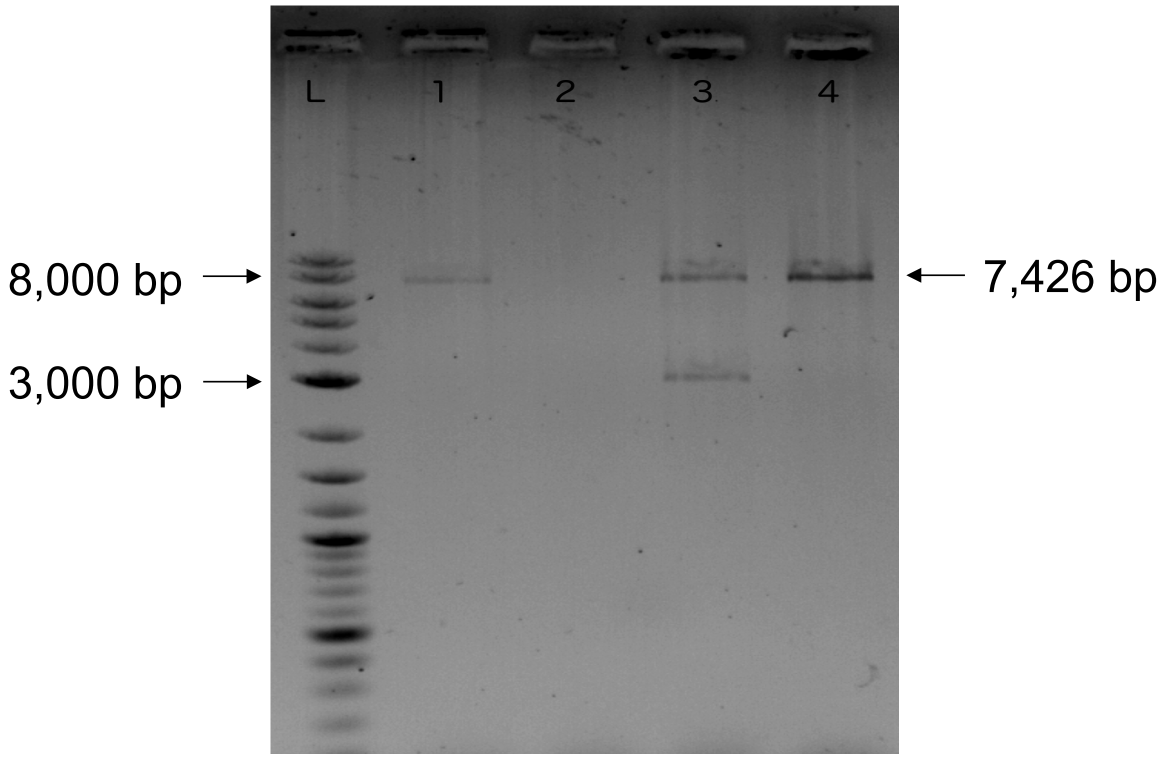


**Figure S9:** PCR amplicons generated from *D. circinale* genomic DNA using primers Acir_sxtG_F and Acir_sxtI_R targeting the region between *sxtG* and *sxtI* in the *sxt* gene cluster.

**References:**


BLIN, K., SHAW, S., STEINKE, K., VILLEBRO, R., ZIEMERT, N., LEE, S. Y., MEDEMA, M. H. & WEBER, T. 2019. antiSMASH 5.0: updates to the secondary metabolite genome mining pipeline. *Nucleic Acids Res,* 47**,** W81-W87.

EL-GEBALI, S., MISTRY, J., BATEMAN, A., EDDY, S. R., LUCIANI, A., POTTER, S. C., QURESHI, M., RICHARDSON, L. J., SALAZAR, G. A., SMART, A., SONNHAMMER, E. L. L., HIRSH, L., PALADIN, L., PIOVESAN, D., TOSATTO, S. C. E. & FINN, R. D. 2019. The Pfam protein families database in 2019. *Nucleic Acids Res,* 47**,** D427-D432.

HAFT, D. H., LOFTUS, B. J., RICHARDSON, D. L., YANG, F., EISEN, J. A., PAULSEN, I. T. & WHITE, O. 2001. TIGRFAMs: a protein family resource for the functional identification of proteins. *Nucleic Acids Res,* 29**,** 41-3.

KANEHISA, M. & GOTO, S. 2000. KEGG: kyoto encyclopedia of genes and genomes. *Nucleic Acids Res,* 28**,** 27-30.

KONSTANTINIDIS, K. T., ROSSELLO-MORA, R. & AMANN, R. 2017. Uncultivated microbes in need of their own taxonomy. *ISME J,* 11**,** 2399-2406.

KURTZ, S., PHILLIPPY, A., DELCHER, A. L., SMOOT, M., SHUMWAY, M., ANTONESCU, C. & SALZBERG, S. L. 2004. Versatile and open software for comparing large genomes. *Genome Biol,* 5**,** R12.

MEDEMA, M. H., KOTTMANN, R., YILMAZ, P., CUMMINGS, M., BIGGINS, J. B., BLIN, K., DE BRUIJN, I., CHOOI, Y. H., CLAESEN, J., COATES, R. C., CRUZ-MORALES, P., DUDDELA, S., DUSTERHUS, S., EDWARDS, D. J., FEWER, D. P., GARG, N., GEIGER, C., GOMEZ-ESCRIBANO, J. P., GREULE, A., HADJITHOMAS, M., HAINES, A. S., HELFRICH, E. J., HILLWIG, M. L., ISHIDA, K., JONES, A. C., JONES, C. S., JUNGMANN, K., KEGLER, C., KIM, H. U., KOTTER, P., KRUG, D., MASSCHELEIN, J., MELNIK, A. V., MANTOVANI, S. M., MONROE, E. A., MOORE, M., MOSS, N., NUTZMANN, H. W., PAN, G., PATI, A., PETRAS, D., REEN, F. J., ROSCONI, F., RUI, Z., TIAN, Z., TOBIAS, N. J., TSUNEMATSU, Y., WIEMANN, P., WYCKOFF, E., YAN, X., YIM, G., YU, F., XIE, Y., AIGLE, B., APEL, A. K., BALIBAR, C. J., BALSKUS, E. P.,

BARONA-GOMEZ, F., BECHTHOLD, A., BODE, H. B., BORRISS, R., BRADY, S. F., BRAKHAGE, A. A., CAFFREY, P., CHENG, Y. Q., CLARDY, J., COX, R. J., DE MOT, R., DONADIO, S., DONIA, M. S., VAN DER DONK, W. A., DORRESTEIN, P. C., DOYLE, S., DRIESSEN, A. J., EHLING-SCHULZ, M., ENTIAN, K. D., FISCHBACH, M. A., GERWICK, L., GERWICK, W. H., GROSS, H., GUST, B., HERTWECK, C., HOFTE, M., JENSEN, S. E., JU, J., KATZ, L., KAYSSER, L., KLASSEN, J. L., KELLER, N. P., KORMANEC, J., KUIPERS, O. P., KUZUYAMA, T., KYRPIDES, N. C., KWON, H. J., LAUTRU, S., LAVIGNE, R., LEE, C. Y., LINQUAN, B., LIU, X., LIU, W., et al. 2015. Minimum Information about a Biosynthetic Gene cluster. *Nat Chem Biol,* 11**,** 625-31.

TATUSOV, R. L., GALPERIN, M. Y., NATALE, D. A. & KOONIN, E. V. 2000. The COG database: a tool for genome-scale analysis of protein functions and evolution. *Nucleic Acids Res,* 28**,** 33-6.
